# Molecular-level observation of the self-assembly of a virus-like particle

**DOI:** 10.64898/2026.02.03.703079

**Authors:** Roi Asor, Dan Loewenthal, Diana Melnyk, Tiong Kit Tan, Philipp Kukura

## Abstract

Biomolecular assembly is a cornerstone of cellular organisation and function. Revealing its underlying principles is essential for understanding biological processes, and their malfunction in disease. Viral capsid assembly is the archetypal self-assembly system that has been central in establishing the fundamental principles of biological self-assembly, providing a conceptual and geometric framework that underpins the current understanding of supramolecular biomolecular systems, the development of new biomaterials, and advancing therapeutic design. Yet, despite decades of experimental efforts, observation and quantification of virus self-assembly pathways and dynamics have remained elusive. Here, we combine mass photometry with a non-perturbative single molecule trapping method, enabling direct, real-time monitoring of the self-assembly of individual virus-like particles (VLPs) with molecular resolution. We show that weak, diffusion-limited, and reversible multivalent interactions control the assembly process by facilitating stochastic selection of a limited set of on-path, topologically closed intermediates. Assembly is finely tuned by the transition rates between these topologically closed configurations and proceeds through a sequence of effectively irreversible first-passage events. The intrinsic first-passage times are a consequence of VLP symmetry, creating a separation of timescales between the formation of the first closed intermediate and subsequent elongation. This separation results in a nucleation and growth mechanism that yields an equilibrium distribution consistent with the law of mass action, despite the overall irreversibility of assembly. Our approach enables direct and complete characterisation of both the thermodynamics and the kinetics governing VLP assembly and reveals how the system achieves specific assembly of one final structure with high fidelity despite the availability of thousands of assembly intermediates. More broadly, our approach provides a general framework for visualising and quantifying the dynamics of multimeric biological machines at the molecular level.

---

Protein self-assembly is crucial for biological function, enabling emergent properties such as structural stability, cooperativity, and dynamic response that proteins cannot achieve individually^1-3^. Driven by a set of non-covalent homo- and hetero-interaction principles^3,4^, self-assembly spans multiple scales and dimensions^5^, from soluble oligomers to surface-associated clusters on membranes. In solution, it underpins the dynamic organisation of the cytoskeleton, the formation of bacterial microcompartments, viral capsids, and membraneless organelles^6-9^, while on membranes it mediates signalling, immune recognition, and virus–host interactions^10-14^. Studying and quantifying assembly across spatial and temporal scales is thus essential for elucidating how structure, symmetry, and dynamics are coupled to enable biological function, how they are dysregulated in disease^15,16^, and how they may be harnessed for therapeutic intervention.

An archetypal example for biomolecular self-assembly process are simple spherical viruses, where identical protein subunits interact to form multivalent building blocks that further assemble into highly regular symmetrical capsids, encapsulating the viral genetic material^17-20^. This process is highly efficient, showcasing self-assembly dynamics finely tuned to distinct phases of the viral life cycle^21^. Acquiring pathway-level understanding of the underlying molecular dynamics is critical for the rational design of new artificial capsid-like protein nanocages^22-24^, as well as the design and development of antiviral treatments^25,26^. In human immunodeficiency virus type 1 (HIV-1), for example, capsid-targeting inhibitors modulate specific pairwise interactions, perturbing both assembly kinetics and the thermodynamic stability of capsids, affecting multiple stages during viral replication^27,28^. In Hepatitis B, capsid assembly modulators bias assembly towards trajectories that lead to malformed or hyperstable particles, suppressing correct nucleic acid encapsidation^29-31^.

The fundamental challenge to visualise the molecular details of capsid assembly comes down to two factors. First, the self-limited free energy landscape involves weak and reversible pairwise interactions^32-35^, meaning that assembly is stochastic, while key intermediates are short-lived and of very low abundance. Second, the involvement of 10s to 1000s of subunits results in an exponentially increasing number of molecular assembly paths and a high degree of heterogeneity in solution^36,37^. As a result, ensemble-based approaches have struggled with this heterogeneity, relying on complex analysis or modelling^31,32,36,38-40^. Single-molecule methods have been limited to ensemble-averaged snapshot observations of stable intermediates due to a lack of temporal resolution^41-44^, a lack of a direct readout of the molecular oligomeric state^45,46,47^, or limited to 2D assembly on a substrate^48-50^. As a result, despite decades of dedicated study, no experimental approach has been successful in visualising capsid assembly^21^. Thus, while theoretical and computational models for assembly are established^51-56^, fundamental questions remain, such as how assembly navigates the highly heterogeneous configurational space, what molecular mechanisms lead to or avoid kinetic traps, and what sequence of molecular events result in capsid polymorphism.^57-60^

Overcoming these limitations requires a new experimental framework. Such a framework must be able to: (i) Quantify the thermodynamic and kinetic parameters that govern the fundamental, pairwise interactions between multivalent subunits. (ii) Monitor particle assembly dynamics at the ensemble level. (iii) Identify transient intermediates and map their relative abundances as a function of time. (iv) Monitor the assembly progression of single complexes by capturing the sequence of microscopic states and the transition rates between them. Here, we establish such a framework by combining solution mass photometry (MP)^61^ with a non-perturbative single molecule trapping approach, enabling complete, molecular-level characterisation of the assembly process of virus-like particles (VLPs).

## The assembly process is gated by weak, μM affinity, pairwise interactions

To enable an analytical understanding of the stoichiometries and topologies involved in the assembly pathway, we selected an engineered, self-assembling virus-like particle (mi3-VLP^62,63^, also known as SC003-mi3), currently under evaluation as a multivalent vaccine platform^25,64-66^. The mi3-VLP was derived from i3-01^62^ by introducing 2 mutations that remove surface-exposed cysteines to prevent aggregation, and by genetically fusing a SpyCatcher domain to the N-terminus to enable plug-and-play attachment of SpyTagged proteins^63^. The mi3-VLP consists of 60 identical mi3-monomers, hierarchically assembled into a dodecahedron from 20 homotrimeric building blocks, yielding a particle with 12 pentagonal faces, sharing molecular similarity with icosahedral native viral capsids (**Fig. 1a**). Although the geometry is dodecahedral rather than the triangulated icosahedral lattice of native viral capsids, the fundamental biophysical principles governing the assembly process are conserved, where symmetry governs the specific pattern of locally stable intermediates (**Fig. 1a**), the degeneracy of microstates, and how assembly pathways branch^34,40^.

**Fig. 1:**
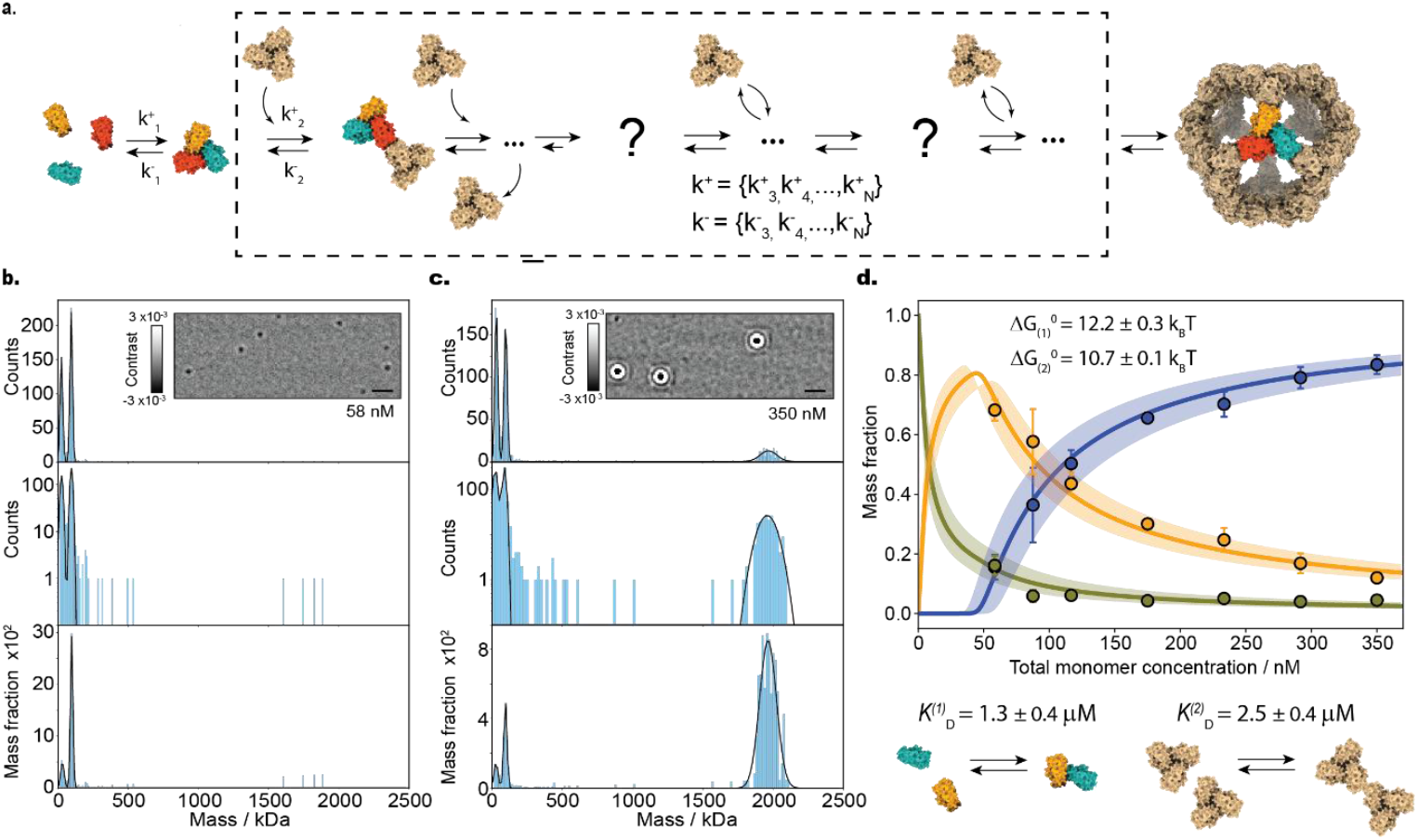
Equilibrium characterisation of the self-assembly process. (a) Illustration of the assembly process for the mi3-based dodecahedral VLP. (b, c) Equilibrium solution mass distributions at two different mi3-monomer concentrations. Insets show representative mass photometry ratiometric frames. Scale bars: 1 µm. Mass histograms are shown on a linear (top panel), semilogarithmic (middle) and mass fractions scale (bottom). Black lines correspond to the sum of three fitted Gaussian functions for the monomer, trimer and VLP mass peaks. (d) Experimental mass fractions for the monomer (green), trimer (orange) and assembled capsid (blue) as a function of the total protein concentration. Symbols and error bars correspond to the average and standard deviation of three technical repeats per concentration. Solid lines correspond to a global fit (**Supplementary note 2.4**) varying the standard free energy change for pairwise interactions between monomers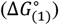 and trimers 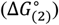.

We begin with a characterisation of the assembly process in solution using standard mass photometry. At 58 nM total mi3-monomer concentration, the solution distribution is dominated by monomers and trimers at 38 ± 2 kDa and 108 ± 1 kDa, respectively (**Fig. 1b**, top and **Figs. S1-S2**). Plotting the mass distribution on a logarithmic scale (**Fig. 1b**, middle) reveals the detection of few intermediate species <500 kDa, and near the mass of the fully assembled VLP (2030 ± 10 kDa). Conversion into mass fractions (**Fig. 1b**, bottom) shows that the mi3-monomers are predominantly found in trimeric form (65 ± 4%) followed by fully assembled VLP (15 ± 4%), monomers (11 ± 3%) and small oligomers (9 ± 3%) (**Fig. 1b**, bottom). At a higher total monomer concentration (350 nM), the majority of protein is now found in assembled VLPs (82 ± 5%) (**Fig. 1c**). We can convert these relative abundances of the three dominant molecular states (monomer, trimer and VLP) as a function of total protein concentration (**Fig. 1d**) into the key affinities governing the assembly process (see **Supplementary note 2.4**),^36,54^ namely 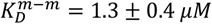 and 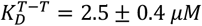 (**Fig. 1d**, bottom). The key pairwise interactions are thus weak relative to the pseudo-critical concentrations needed for assembly (∼50 nM), which emphasises the importance of multivalency in stabilising these interactions.

## Reduced dimensionality from 3D to 2D promotes pentagonal face assembly

A major challenge for characterising self-limiting assembly in solution is the extremely low abundance of assembly intermediates^35,67,68^. In our case, the intermediate states are those originating from the assembly of trimers into larger oligomers, on-path to the full VLP. One approach to make higher order assembly more favourable is to reduce the entropic penalty associated with complex formation. We achieve this here by tethering histidine-tagged trimeric subunits to supported lipid bilayers where they are able to freely diffuse in two dimensions (**Supplementary note 3**). In this 2D configuration, rotational degrees of freedom are constrained, pre-aligning trimers for oligomerisation, increasing the interaction free energy gain^69,70^, and increasing the effective concentration (**Fig. 2a**). Since mi3-subunits do not interact with the SLB in the absence of a tag, we expect tethering to the 2D surface to maintain the solution structure and conformation.

**Fig. 2:**
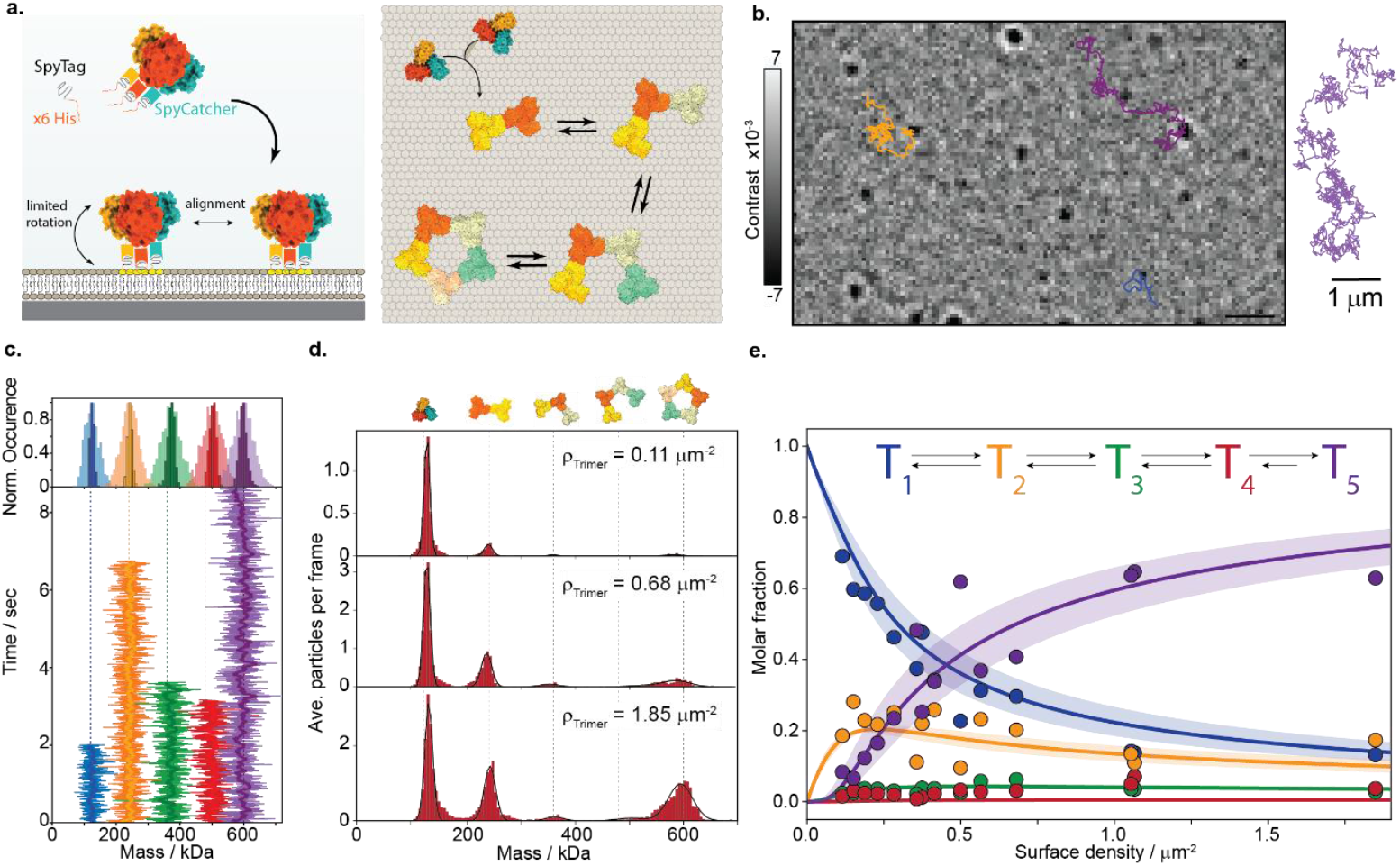
Dimensionality reduction induces the assembly of pentamers of trimers on supported lipid bilayer. (a) Tethering of mi3-subunits to a supported lipid bilayer using polyhistidine NTA-Ni lipids linking. (b) On the SLB, individual proteins and their complexes are imaged, tracked and mass monitored as they diffuse and self-assemble on the SLB (Dynamic-MP). Left: Representative MP image acquired at 270 Hz (**Supplementary notes 3-4**). Representative partial trajectories are shown for a trimer (blue), dimer of trimers (orange) and pentamer of trimers (purple). Scale bar: 1 µm. Right: An 8 second trace of a diffusing pentamer. (c) Representative mass traces of different oligomeric states, measured at the raw instrument frame rate (270 Hz), and after 10 frame averaging (37 ms). (d) Surface mass distributions as a function of trimer surface density. (**Supplementary notes 4.3-4.5**). Vertical lines and illustrations show the expected masses of the different oligomers of trimers. Each histogram contains cumulative data from between 3 (for high density) and 15 (for lower density) MP measurements. Black lines correspond to the sum of the best fitted Gaussian functions used for fitting each mass peak. (e) Mole fractions as a function of trimer surface density (symbols) and corresponding thermodynamic fit (lines) (**Supplementary note 5**).

In contrast to the experiments in **Fig. 1** where we characterise solution distributions from non-specific binding of proteins to a glass surface, the data now consists of individual proteins diffusing in 2D on an SLB^71-73^ (**Fig. 2b, Supplementary note 4**). Different oligomers of trimers appear at their respective mass multiples (**Fig. 2c**), and are resolvable by mass even at the raw frame rate of our measurement (270 Hz). In analogy to a solution-based experiment, where increasing the solution concentration leads to a higher fraction of assembled capsids (**Fig. 1c**), raising the surface density increases the abundance of higher-order oligomers (**Fig. 2d**). We observe the expected exponential decay in higher oligomer abundance with the exception of the pentamer of trimers, which is stabilised by the additional free energy associated with ring formation. Larger assemblies are not formed since the mi3-subunits solution concentration is well below the pseudocritical concentration for assembly (<5 nM), while surface tethering prevents out of plane interactions. In this way, we arrest the assembly and characterise the formation of the smallest stable intermediate, a pentagonal face.

Resolving the complete oligomeric distribution while directly measuring the trimer surface density allows us to construct a complete thermodynamic model for 2D assembly. Here, we require only one fitting parameter, the pairwise, standard 2D association free energy change, Δ*G*^∘^(2*D*) (full description of the model in **Supplementary note 5**). Fitting the oligomeric occupancies as a function of surface density (**Fig. 2e**) yields a 2D interaction free energy gain of 6.75 ± 0.2 *K*_*B*_*T* per contact that corresponds to 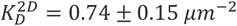,for trimeric subunits.

We can now use the ratio of the 2D and 3D dissociation constants, which defines a characteristic confinement length 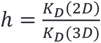, to reveal the origin of affinity enhancement upon surface confinement. In our case, *h* = 0.5 ± 0.1 *nm*, falling within the estimated values for enhancement arising from restricted translational and rotational degrees of freedom in two dimensions^70^. By contrast, a substantial membrane-induced enthalpic stabilisation would result in values orders of magnitude smaller. Therefore, we conclude that, thermodynamically, our surface-based measurement is representative of measurements in solution, the only exception being a different scaling originating purely from the reduction in dimensionality.

Thermodynamic analysis (**Figs. 1d and 2e**) suggests that the formation of this pentagonal face on the SLB, and the formation of the complete VLP in solution, are consistent with a single trimer–trimer contact free energy. Accordingly, we expect that any transiently stabilised intermediates or variability in transition times between states are unlikely to arise from cooperativity or alternative conformations, which would result in different fundamental interaction parameters. Instead, any assembly heterogeneity is topology driven: as assembly proceeds, the local coordination (number of intertrimer contacts per subunit) increases, and ring closure adds further stabilisation, thereby governing intermediate lifetimes and transition kinetics.

## Pairwise association of trimers is diffusion-limited

SLB-tethering of subunits enables continuous monitoring of the molecular masses, positions, mobilities and interactions between individual subunits (**Fig. 3a-b**), a measurement that cannot be performed in solution due to rapid diffusion. Thereby, we can quantify the molecular dynamics directly, extending the thermodynamic characterisation. In the language of chemical reaction dynamics, imaging the relative positions of particles yields reaction coordinates in terms of the approaching subunits, while the measured mass reveals the oligomeric state and any associated changes, analogous to the making and breaking of chemical bonds. In this way, we can track the molecular states as a function of time and characterise the time spent in each state, the dwell time, and any transitions to a different state (**Fig. 3a**).

**Fig. 3:**
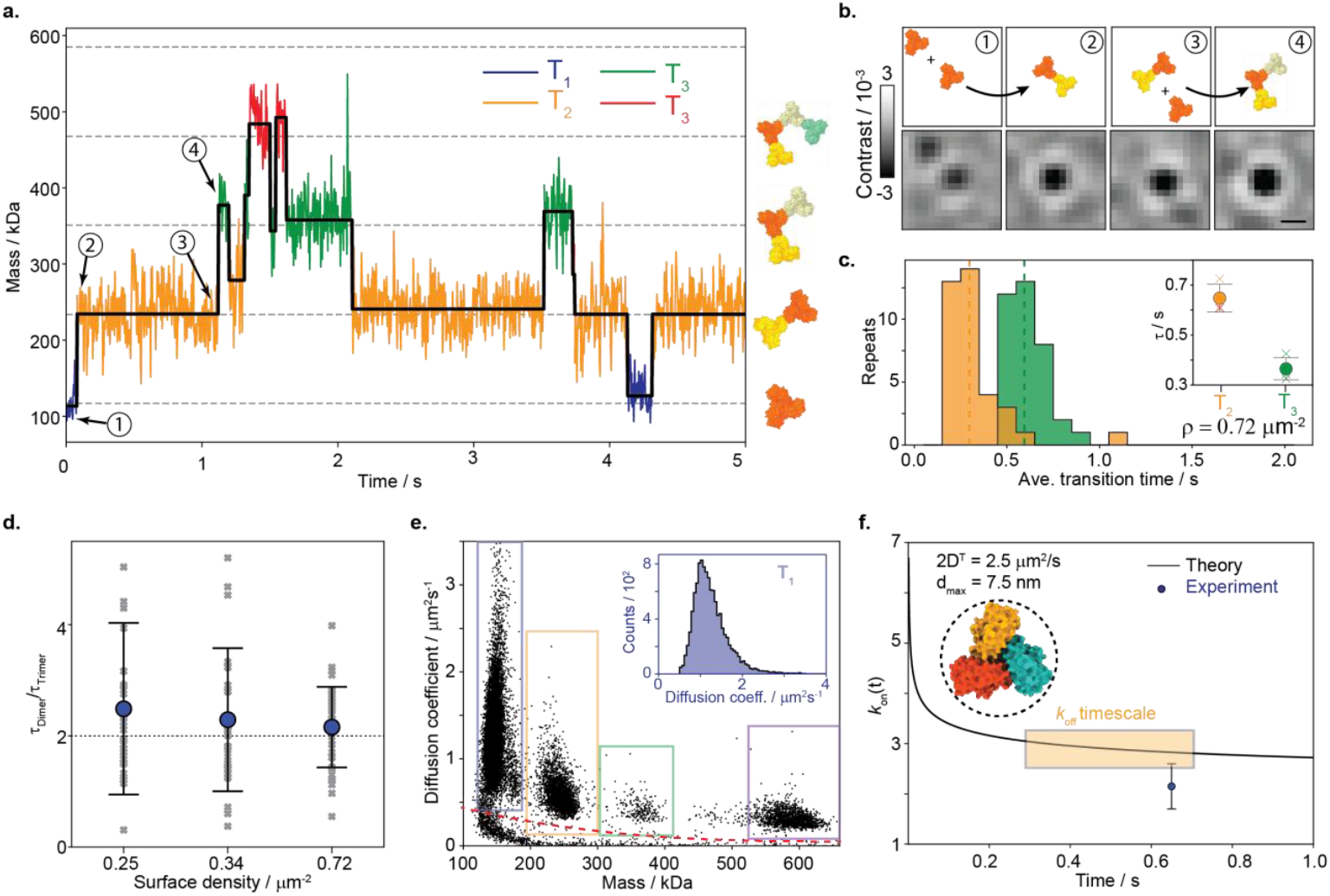
Diffusion limited pairwise interactions guide the assembly of the pentamer of trimers. (a) Continuous mass monitoring during individual trajectories (coloured, see **Movie S1**) and corresponding outcome of a step-finding algorithm (black curve, see **Supplementary note 4**). (b) Zoom of dimer and trimer formation. Scale bar: 300 nm. (c) Representative average transition time histograms for dimer (orange) and trimer (green) disassembly. The average values for each distribution are indicated by a vertical line. Inset: Circles indicate the averaged values for the dimer (orange) and the trimer (green) over different surface densities, where crosses indicate the values at the three different surface densities, and the error bars represent the standard deviations. (d) The ratio between the averaged transition time of the dimer with respect to the trimer within the same measured movie, calculated at different surface densities. Crosses indicate the measured ratio within individual movies, full circles and error bars represent the averages and the standard deviations, respectively. (e) The molecular mass and the diffusion coefficient measured for individual oligomers (scattered symbols). The area below the red dashed line indicates noise contributions from the SLB. Inset: the diffusion coefficient distribution measured for trimeric subunits. (f) The mobility information together with the calculated dimensions of the molecular species (PDB ID: 7b3y) yields the theoretical, diffusion-limited, two-dimensional forward reaction rate constant, 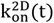 for the formation of dimer of trimers (**Supplementary note 6.5**).

The resulting transition rates are 0.65 ± 0.05 s for the dissociation of dimers of trimers and 0.36 ± 0.04 s for dissociation of trimers of trimers (**Fig 3c** and **Fig. S4**), yielding *K*_*off*_= 1.5 ± 0.1 *s*^−1^ and 2.8 ± 0.3 *s*^−1^, respectively (see **Supplementary note 6**). Combination with our 2D thermodynamic characterisation yields the complete reaction dynamics for formation of the pentagonal face. The bimolecular *K*_*on*_ for dimerisation is given by 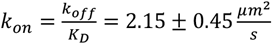 and the same calculation for the trimer formation yields 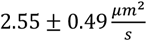. To place these values in context, we estimate the 2D diffusion-limited rate constant 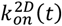^74^ using our experimental quantification of the 2D diffusion coefficient per particle, which is inversely proportional to the oligomeric state (**Fig. 3e** and **Supplementary note 6.5**). Focusing on dimerisation, the subunits have an average diffusion coefficient of 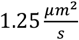 (**Fig. 3f**, top) and an encounter diameter of 7.5 nm (estimated from the atomic structure^75^). Evaluated between 0.3 and 0.7 s, which is the timescale relevant to the observed dissociation, yields 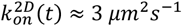 (**Fig. 3f**, bottom), in good agreement with our experimentally determined value. These results show that association between the trimeric subunits is essentially diffusion-limited with no appreciable energetic barriers for association, where sampling the correct orientation between the two trimeric subunits is likely the limiting factor.

## Assembly proceeds via topologically stable intermediates

By separating the assembly process into two phases, solution and SLB (**Fig. 4a**), we can create a modest driving force for assembly, while keeping the assembly conditions close to equilibrium. We achieve this by producing an equilibrated surface mixture of oligomers that contains closed pentameric rings, which serve as a nucleus for assembly. They represent the smallest stable oligomeric state with a higher probability to elongate than dissociate. In solution, untagged mi3-subunits form an equilibrated mixture where mi3-monomers, trimers and complete VLPs coexist (**Fig.1d**). Since each phase by itself is equilibrated, there is no net driving force for assembly within each phase. However, coupling the equilibrated systems by bringing preassembled surface pentamers into contact with the equilibrated solution mixture (containing the molecular species shown in **Fig. 1**), introduces a chemical potential difference between the pentagonal ring and the complete VLP, mediated by the pseudo-critical concentration of free trimeric subunits. In this coupled configuration, supersaturation (**Supplementary note 8**) between surface pentamers and complete VLPs is constant, because the bulk concentration is effectively unchanged.

**Fig. 4:**
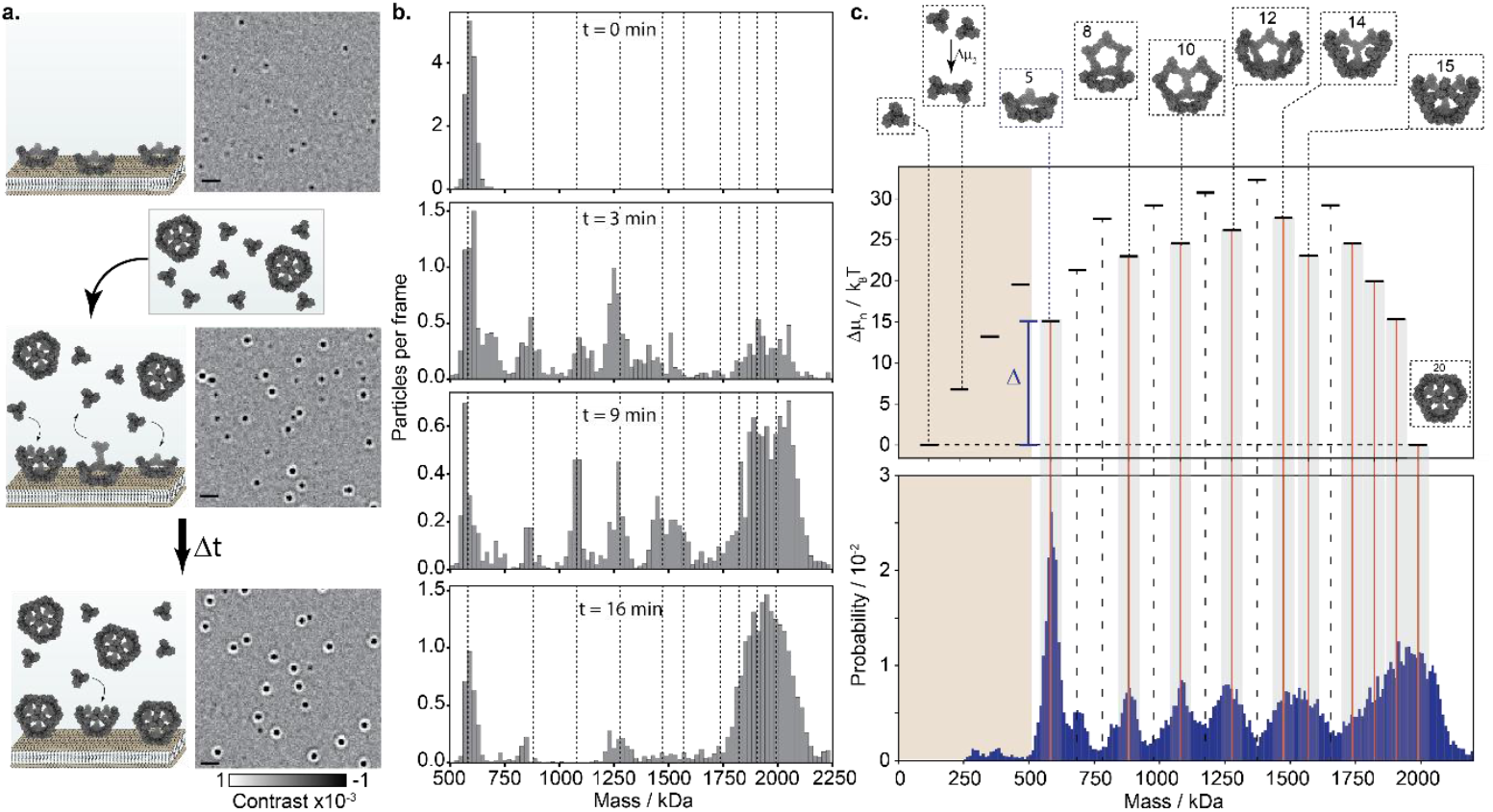
Stable intermediates reveal the grand canonical free energy landscape of VLP assembly. (a) Driven VLP assembly on SLBs. Surface-assembled pentameric rings (top) are put in contact with equilibrated trimers and capsids in solution (middle), which leads to capsid assembly on the SLB (bottom). Scale bar: 1 µm. (b) Resulting mass histograms as a function of time after addition of 44 nM mi3 protein in solution. Vertical lines correspond to the masses of closed-structure intermediates. Mass broadening, owing to particle size, for masses >1750 kDa prevents resolving individual molecular states at the ensemble level. (c) Time-averaged mass distribution (between 4 and 15 minutes) and comparison with the calculated grand-canonical free energy landscape using interaction parameters from **Fig. 1**. Solid red lines: Topologically closed structures based on PDB ID: 7b3y. Dashed lines: Unstable configurations.

The resulting small driving force (∼15*K*_*B*_*T*), together with diffusion-limited and reversible inter-trimer interactions, results in a slow assembly reaction, enabling near-equilibrium sampling of the assembly propagation pathway. Importantly, subunits in solution lack the histidine tag and therefore do not interact with the SLB (**Fig. S7**). Any observed intermediates thus arise from tethered pentamers that increase in mass, and not from SLB-adsorption of particles from solution (**Fig. 4b**). Repeating the experiment at decreasing solution concentration slows down the assembly timescale, ranging from ∼10 min for complete assembly at 88 nM, ∼20 min at 44 nM (**Fig. 4b**) and 40 min at 29 nM (**Figures S9-S16**). Although the timescales differ, the involved intermediate states (indicated in **Fig. 4b**, black vertical lines) are similar across different concentrations. Averaging distributions across technical repeats during time periods where relatively little change in intermediates is observed (**Fig. S18**) improves the statistics and reveals well-resolved peaks and valleys as a function of mass.

Overlaying the calculated free energy landscape (**Supplementary note 8** and **Fig. S17**) of the lowest free energy path between the trimeric subunit and the complete VLP reveals the driving force for assembly following the formation of the pentagonal ring (**Fig. 4c**, top). It also shows that mass states that accumulate along the assembly reaction match local minima in the free energy landscape, while valleys correspond to local maxima (**Fig. 4c and Fig. S17**). Examining the associated structures shows that stable intermediates are those composed of closed pentagonal faces (or closed rings), stabilised by a larger trimer-trimer coordination number. It is important to note that the equilibrated grand canonical free energy landscape, representing the equilibrated solution, does not change when the two systems are coupled, only the starting point for assembly changes. Therefore, the measured distribution of oligomeric species on the bilayer propagates in time according to the molecular transitions governed by the equilibrated underlying free energy potential.

## Single particle dynamics reveal irreversible step-like transitions between topologically-closed intermediates

Population-level measurements highlight overrepresented states and thus hint at key assembly intermediates. From an ergodic standpoint, repeated measurements of one complex should converge to the ensemble probability distribution. However, the ensemble does not provide information on the sequence of steps, or the transition dynamics. We therefore developed an experimental approach based on mass photometry to observe the real-time assembly dynamics of VLPs at the level of individual complexes with molecular resolution. To achieve this, we create circular SLBs of 5 µm diameter by lithographic removal of a dense poly-L-Lysine polyethylene glycol (PLL-PEG, **Fig. 5a**, top and **Fig. S20**) layer.^76^ Any species bound to the SLB are now confined, and can thus be observed and mass measured for extended periods of time (**Supplementary note 9**).

**Fig. 5:**
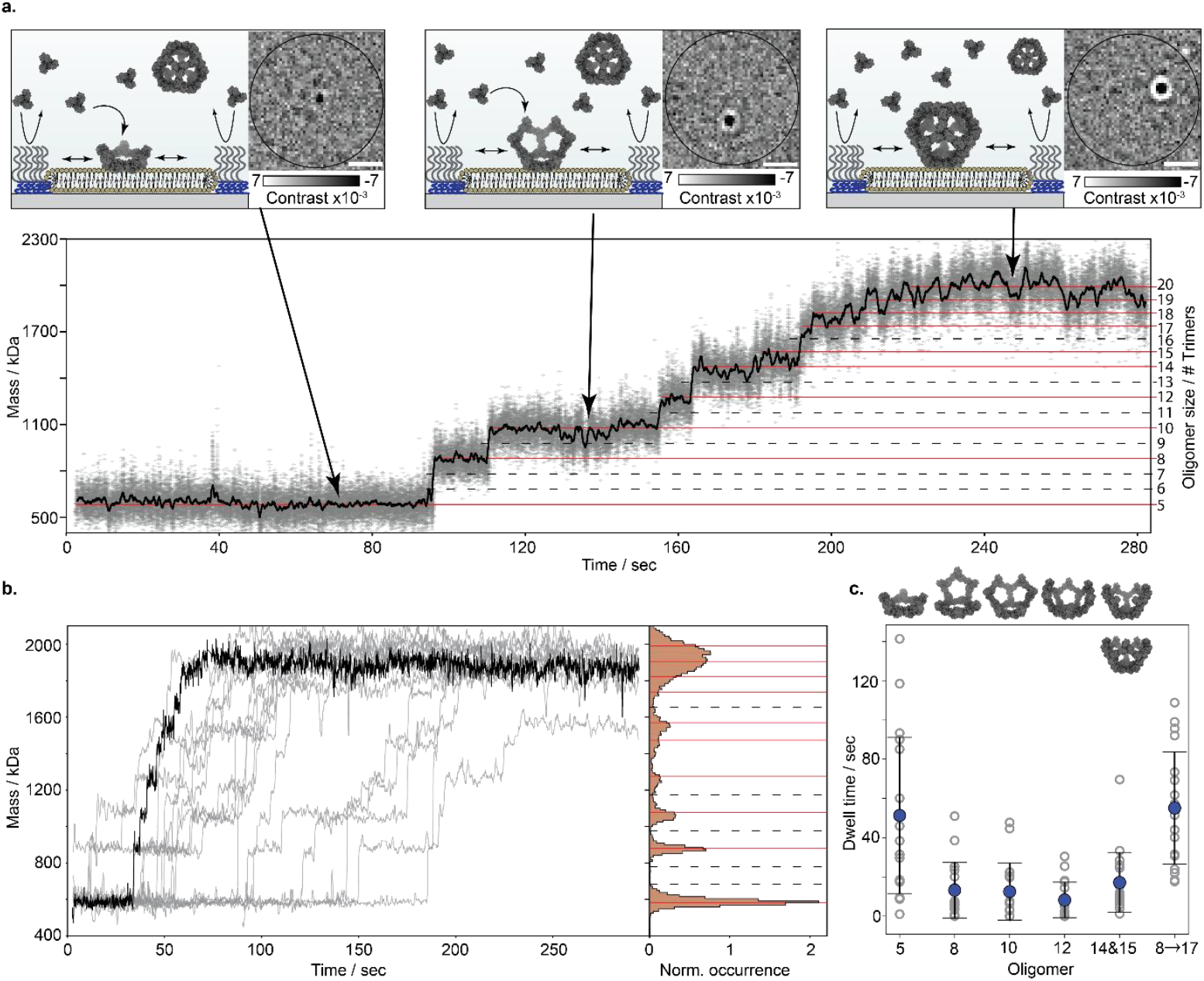
Single-molecule observation of self-assembly pathways and dynamics. (a) Top: Illustration of single complex mass photometry by SLB traps. Scale bar: 1 µm. Bottom: Mass trace for the assembly of a single VLP. Grey symbols show the measured mass at 250 Hz imaging frame rate (**Movie S2**), and the black line a running median with a symmetric window size of 100 frames (400 ms). Red (stable) and black (unstable) horizontal lines denote the expected masses of complexes along the lowest free energy path from the pentameric state to the complete assembled VLP (**Fig. 4a**). (b) 18 superimposed mass traces capturing the assembly of individual VLPs. Traces are shown following the application of a running median with a window size of 100 frames (400 ms). The histogram to the right corresponds to the average normalised distribution of the sampled mass states throughout the presented trajectories. Horizontal lines correspond the same states as in (a). (c) Calculated dwell times for the most stable intermediates along the assembly path, and the time interval between the formation of the first intermediate structure (8-mer) and the formation of a particle that is equal or larger than 17 subunits (representing the transition through all intermediate states for the assembly process). The data was obtained from the 18 single-molecule trajectories in (b) (**Supplementary note 8.4 and Fig. S21**). Grey symbols indicate the individual values from each trajectory, where full blue symbols and error bars represent the averages and standard deviations.

We form SLB-bound pentameric rings by adding his-tagged mi3-subunits as previously but now to SLB traps, followed by ∼30 min of equilibration. Subsequently, we add an equilibrated solution of unlabelled mi3-subunits at total protein concentrations between 88 and 100 nM. Trimeric subunits from solution now bind to tethered pentameric rings, which we monitor in real time by continuous mass measurement (**Fig. 5a** and **Movie S2**). The resulting mass traces reveal the oligomeric states that are populated for extended time periods, separated by sharp mass changes. Intermediate masses match those of topologically closed structures (red lines), that also accumulate in ensemble experiments (**Fig. 4d**). We observed these features irrespective of assembly rate, where both slower (**Fig. 5a**, top) or faster (**Fig. 5b**, black curve) molecular mass trajectories exhibited similar features.

Superimposing multiple trajectories (**Fig. 5b**) confirms these observations, where the normalised distribution exhibits features similar to the equilibrium results (**Fig. 4c**), linking the single complex dynamics experiments (**Fig. 5b**) to population mass distributions (**Fig. 4c**). While the rapid off-rate combined with the small mass change (108 kDa) associated with pairwise trimer-trimer interactions (**Fig. 1**) is beyond the capabilities of our current experimental approach, we can observe and identify the key states with longer dwell times, i.e. those representing closed structures. Notably, we do not observe disassembly in any of the measured trajectories.

When evaluating the lifetimes associated with different intermediates, the pentameric ring exhibited the longest dwell time, while other species showed broadly similar dwell times (**Figure 5c**). This is readily explained from a molecular perspective: Transitioning from the pentagonal ring to the next stable state (8-mer) requires binding of three weakly interacting subunits to form an additional topological face before any of them dissociates. The subunit concentration in solution determines the collision frequency, and therefore the probability that three subunits bind in the correct configuration prior to unbinding. By contrast, for subsequent transitions up to the 14-mer, closing each new face requires only two additional subunits, leading to shorter transition times. We note that the reported pentamer dwell time is a lower bound due to the ∼20 s gap between subunit addition and the start of acquisition. Because the 14 and 15-mers are both closed structures, we report the cumulative dwell time (top of **Fig, 5c**).

## Discussion

Our measurements reveal the molecular, energetic and kinetic details of a topology-driven, self-limiting, VLP assembly pathway (**Fig. 6a**). Trimeric subunits engage in weak pairwise interactions (*K*_*D*_∼2*μM*) allowing for reversible encounters, which stochastically explore molecular configurations until a topologically-closed stable arrangement is reached. Importantly, the short lifetime of individual interactions prevents the persistence of loosely connected structures. Once a subunit attains a coordination number ≥ 2, the associated stabilisation results in a much slower off-rate than the on-rate for new subunits arriving. Repetition of this process throughout the growth of the dodecahedron enables an efficient and well-defined assembly process.

**Fig. 6:**
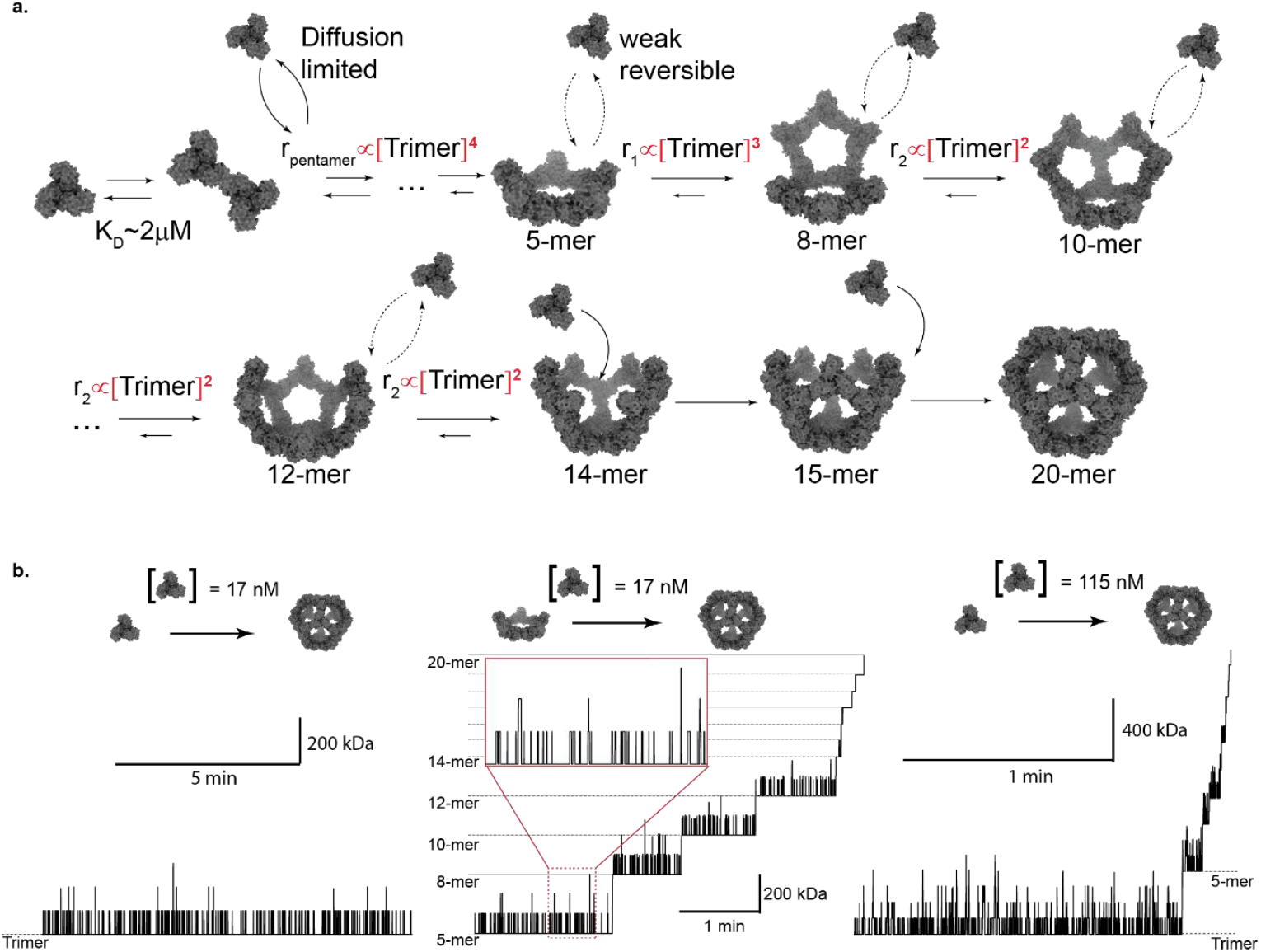
Molecular mechanism of capsid assembly. (a) Quantitative illustration of the assembly mechanism based on the combined bulk and single molecule results. (b) Stochastic simulation of the assembly process assuming the parameters *K*_*on*_ = 10^7^ *M*^−1^*s*^−1^ and *K*_*D*_ = 1*μM*. Left: representative simulated trace starting from a trimeric subunit, at trimer solution concentration of 17 nM. Middle: Starting from a pentagonal face individual subunits from a solution containing 17 nM of trimers bind weakly and reversibly until they randomly bind in a configuration that allows the formation of a ring structure that cannot disassemble. The assembly process proceeds through these stable states. Right: Kinetics of VLPs assembly with the same molecular parameters. This time the starting point is a trimeric subunit and the concentration in solution is 115 nM.

Our results provide a molecular level explanation to the kinetic origin of the difference between nucleation and elongation, the hallmark of a two-states assembly process (**Fig. 1**) and characteristic of many capsids and VLPs^35,68^. The smallest stable intermediate is a pentagonal ring, formed after four trimer additions. The subsequent transition (5-mer to 8-mer) requires three additions, while later steps up to 14-mer requiring only two. The corresponding reaction rates thus scale with the fourth, third and second powers of the trimer concentration, respectively (**Fig 6a**). Due to the irreversibility of the closed topologies, transitions between these configurations can be viewed as a series of first passage stochastic events. Because the first-passage time is highly sensitive to the number of additions, formation of the first pentagonal face is the critical step: Once formed, it enables propagation of the entire assembly process, with sequentially-increasing rates. The distinction between nucleation and elongation thus arises directly from the difference in the number of subunits required to complete topological rings, corresponding to a sequence of first-passage events.

Observation of single capsid assembly events shows that, within the timescale of assembly, the reaction proceeds through a sequence of irreversible, specific steps. This creates an apparent contradiction between microscopic irreversibility that predicts total consumption of trimeric subunits, while our solution data (**Fig. 1**) shows coexistence between subunits and complete VLPs, consistent with the law of mass action^77,78^. Key here is that although specific states are irreversible, the pairwise interactions leading to these states are weak and reversible. Thus, as the assembly reaction proceeds and the concentration of free subunits decreases, the timescale for creating the pentameric intermediate rapidly increases. At the pseudocritical monomer solution concentration, which is eventually reached independent of the starting concentration as capsids are assembled, this timescale becomes either so long that assembly is practically irrelevant, or comparable to the timescale of ring disassembly (longer than our observation window). In both cases, experimentally, the result complies with the expected observation of the law of mass action where subunits and complete VLPs coexist in solution.

We can now understand the assembly process in full detail, represented by a simple stochastic kinetic simulation (**Supplementary note 10**). At sufficiently low concentrations, lower or equal to the pseudocritical trimer concentration (17 nM, **Fig. 1d**), the probability of 5 trimers coming together before at least one of them departs becomes vanishingly low. As a result, the pentagonal ring is never formed and thus, also no VLPs (**Fig. 6b**, left). If, however, we skip this low probability initial pentagonal ring formation step, here by lowering the barrier using an SLB, assembly does become possible at the same solution concentration (**Fig. 6b**, middle and **Fig. 4**). Finally, at concentrations that result in a high assembly yield, the timescale difference between ring formation and subsequent propagation, makes intermediates so transient that at equilibrium only free subunits and complete VLPs will coexist (**Fig. 6b** right and **Fig. 1b-c**).

Beyond the implications for our understanding of the fundamental principles driving and controlling self-assembly, our approach represents a novel framework for studying biomolecular assembly and dynamics more broadly. The approach of characterising pairwise interactions, quantifying equilibrium distributions and monitoring individual assembly events and dynamics is general: Here, it was applied to a simple, homo-oligomeric, analyte, but there are no fundamental limitations that would prevent our approach to be applied to essentially any multi-molecular system, opening up completely new avenues for studying biomolecular dynamics and mechanisms in the context of biological function and regulation.

## Acknowledgments

We thank Jan Christoph Thiele and Andrew Baldwin for helpful discussions and insightful comments, Mark R, Howarth and Rory A Hills for providing the SC003-mi3-VLPs used for initial experiments, and Franziska L. Sendker, Georg K. A. Hochberg for providing the citrate synthase used for the spatial calibration of the contrast. We thank Seham Helmi and Jack Bardzil for careful reading of the manuscript and for helpful comments. R.A. was supported by EMBO long-term postdoctoral fellowship (ALTF-198-2020) and is supported by the EPSRC (EP/T03419X/1). D.L. is supported by a Clarendon scholarship, a Menasseh Ben Israel scholarship, and a Kingsgate scholarship. T.K.T and DM are supported by a Coalition of Epidemics Preparedness Innovation grant under the broadly protective coronavirus vaccine (BPCV) portfolio. P.K. is supported by the EPSRC (EP/T03419X/1 and EP/W001055/1).

## Author contributions

RA, DL, TKT and PK conceptualized the study, RA, DL, TKT and PK designed the research, RA and DL performed the experiments and analysed the data, RA wrote the analysis software, performed the modelling and simulations, DM and TKT expressed, purified and characterised the mi3-VLPs. RA and PK supervised the research. RA, DL and PK wrote the original draft and all the authors contributed to writing, reviewing and editing the paper.

## Declaration of interests

The authors declare the following competing interests: PK is a non-executive director, shareholder of and consultant to Refeyn Ltd. RA, DL and PK have applied for a patent for confined diffusion mass photometry (N432337GB). The other authors are not aware of any affiliations, memberships, funding, or financial holdings that might be perceived as affecting the objectivity of this manuscript.

## Data and code availability

The raw data and code required to reproduce all of the manuscript figures will be deposited in the University of Oxford Research Archive upon manuscript acceptance.

