## Supplemental Information for "Molecular-level observation of the self-assembly of a virus-like particle"

#### Table of Contents

**Supplementary notes: ... 4**
**Supplementary note 1: Materials ... 4**
1.1 Buffers lipids and commercial proteins ... 4
1.2 Expression and purification of SC003-mi3 VLP ... 4
**Supplementary note 2: Solution characterisation ... 4**
2.1 Solution self-assembly experiments ... 4
2.2 Mass photometry measurements in solution ... 5
2.3 Difference in measured mass of the complete VLP ... 5
2.4 Solution thermodynamic model (Figure 1d) ... 6
**Supplementary note 3: Dynamic MP measurements on supported lipid bilayers ... 6**
3.1 Supported lipid bilayer preparation ... 6
3.2 Preparation of the histidine tag mi3-VLPs for measurements on SLB ... 7
**Supplementary note 4: Dynamic MP acquisition and data analysis (Figures 2-3) ... 7**
4.1 Data acquisition ... 7
4.2 Image analysis ... 7
4.3 Segmenting trajectories using step detection ... 8
4.4 Extraction of oligomeric mass and diffusion coefficient ... 8
4.5 Plotting mass histograms and calculating surface molar fractions ... 9
**Supplementary note 5: Characterising two-dimensional interactions ... 9**
5.1 Thermodynamic characterisation of 2D thermodynamic model for Figure 2d ... 9
5.2 Quantifying of the effect of dimensionality reduction ... 10
**Supplementary note 6 : Measurements of oligomeric dwell times ... 10**
6.1 Measurements and mass changes detection ... 10
6.2 Dynamic MP simulations for testing the extracted lifetimes ... 11
6.3 Simulation procedure ... 11

### Supplementary notes:

#### Supplementary note 1: Materials

##### *1.1 Buffers lipids and commercial proteins*

HEPES (H3375), Chloroform (288306), Magnesium Chloride hexahydrate (M2670), Sodium Chloride (S3014) and Trizma base (T1503) were purchased from Merck Life Science UK Limited. Dulbecco's Phosphate Buffered Saline (DPBS) was purchased from ThermoFisher Scientific. UltraPure 10% SDS (15553-035, Invitrogen). Lipids, 1-palmitoyl-2-oleoyl-glycero-3-phosphocholine, 850457 (POPC); 1,2-dioleoyl-sn-glycero-3-[(N-(5-amino-1-carboxypentyl)iminodiacetic acid)succinyl] (nickel salt), 790404P (DGS-NTA) and 1,2-dioleoyl-sn-glycero-3-phosphoethanolamine-N-[methoxy(polyethyleneglycol)-550] (ammonium salt), 880530 (18:1 PEG550 PE) were purchased from Avanti Polar Lipids. The lipid powders were kept at -80 °C and aliquots were stored in chloroform at −20 °C. SpyTag-polyHistidine peptide was purchased from BioMatik, sequence: RGVPHIVMVDAYKRYKGSGESGSGHHHHHH.

##### *1.2 Expression and purification of SC003-mi3 VLP*

The plasmid pET28a-SpyCatcher003(SC003)-mi3 (gift from Professor Mark Howarth, Cambridge University) was transformed into *E. coli* BL21(DE3) RIPL cells (Agilent) and plated on Luria-Bertani (LB) agar supplemented with 50 µg/mL kanamycin. After incubation for 16 h at 37 °C, a single colony was used to inoculate 10 mL LB media containing 50 µg/mL kanamycin and grown overnight at 37 °C with shaking at 200 rpm. This starter culture was transferred into 1 L LB medium with the same antibiotic and incubated at 37 °C, 200 rpm, until the OD600 reached ~0.6. Protein expression was then induced with 0.42 mM IPTG, and cultures were grown for a further 16 h at 22 °C with shaking (200 rpm). Cells were harvested by centrifugation at 4,000 × g for 15 min. Cell pellets were resuspended in 40 mL lysis buffer (20 mM Tris-HCl, 300 mM NaCl, pH 8.5 at 4°C) containing 0.1 mg/mL lysozyme, cOmplete EDTA-free protease inhibitor cocktail (Roche, 1 mg/mL), and 1 mM PMSF. The suspension was passed through a high-pressure homogeniser (Constant Systems) at 30,000 psi, with 2–3 passes on ice. The lysate was clarified by centrifugation at 35,000 × g for 45 min at 4 °C, and the supernatant was collected. Ammonium sulfate was then added at 170 mg per mL of lysate, and the mixture was incubated at 4 °C for 1 h with agitation (220 rpm) to precipitate the particles. Following centrifugation at 30,000 × g for 35 min at 4 °C, the pellet was resuspended in 8 mL buffer (25 mM Tris-HCl, 150 mM NaCl, pH 8.0 at 4°C) and passed through 0.22 µm filters (Croning). The filtrate was dialysed overnight against a 500-fold excess of the same buffer at 4 °C. Dialysed material was centrifuged at 17,000 × g for 30 min at 4 °C to remove insoluble aggregates and filtered again (0.22 µm). Purification was completed by size-exclusion chromatography on a HiPrep Sephacryl S-500 HR 16/60 column (GE Healthcare) equilibrated in 25 mM Tris-HCl, 150 mM NaCl, pH 8.0 at 4 °C, using an ÄKTA Pure 25 system (GE Healthcare). Elution was performed at 1 mL/min, collecting 1 mL fractions. Fractions containing SpyCatcher003-mi3 nanoparticles were pooled, concentrated and dialysed into TBS 25 mM Tris-HCl, 150 mM NaCl, pH 8.0 at 4°C) using a 100 kDa MWCO centrifugal filter (Millipore), and stored at −80 °C. Final protein concentration was determined by BCA assay (Pierce, ThermoFisher).

#### Supplementary note 2: Solution characterisation

##### *2.1 Solution self-assembly experiments*

A previous study[1] has shown that VLPs can reversibly disassemble and reassemble at guanidinium thiocyanate concentration of 2.5 M. To perform the assembly experiments we started with a solution containing purified VLPs (see **Fig. S1**) at a monomer concentration of 84 $\mu$M, and disassembled the particles using 2.5 M of guanidinium thiocyanate at varying protein concentrations ranging from 35 $\mu M$

to 5.8 $\mu M$ of mi3-monomers. Following an incubation time of 30 to 60 min at room temperature, assembly was initiated by rapidly diluting the protein solution 100-fold in assembly buffer (20 mM HEPES pH 7.4 138 mM NaCl) to a final mi3-monomers concentration ranging between 350 and 58 nM. The assembly reactions equilibrated at room temperature for an additional 30 min. We quantified the distribution of masses of the reassembled VLPs at the chosen concentrations using a standard MP solution assay.

##### *2.2 Mass photometry measurements in solution*

The equilibrated assembly reactions (Fig. 1 and S1-2) were measured using a commercial mass photometer (TwoMP Refeyn Ltd., Oxford) using an imaging field of view of 4.3×10.9 $\mu m^2$. Measurements were conducted on microscope glass coverslips (24 × 50 mm, Menzel Gläser, VWR 630-2603) that were cleaned by three consecutives 5 min cycles of bath sonication in acetone, 50% isopropanol in Milli-Q® water (18.2 MΩ·cm), and Milli-Q. Cleaned coverslips were then dried using nitrogen flow and 3 mm silicone gaskets (GBL103250, Grace Bio-Labs) were attached on the coverslip surface. The gasket was prefilled with 15 µL of buffer and the focus position was adjusted for optimal contrast before adding 5 µL of protein solution. Measurements were performed at a frame rate of 500 Hz followed by frame binning of 2, resulting in an effective frame rate of 250 Hz. Data analysis was performed using DiscoverMP v2024R1 (Refeyn Ltd.) in which rolling ratiometric movies were generated using an averaging window size of 20 frames (80 ms). Threshold parameters for particle detection were set to the default values of 1.5 (threshold 1) and 0.25 (threshold 2). For each data set, a calibration of interferometric contrast to mass was performed using a protein standard while using the same acquisition parameters, similar to a previously reported procedure [2,3].

##### *2.3 Difference in measured mass of the complete VLP*

When examining the measured mass of the soluble mi3-trimers the average measured mass is $107.5 \pm 1.2\ k$Da (**Fig. S1a**). Therefore, the expected measured mass of the mi3-VLP is expected to be $2150 \pm 24\ kDa$. However, the particles are measured at $2031 \pm 10k$Da (**Fig. S1b,** top), slightly below the expected mass. The origin of this small change in the measured interferometric contrast originates from the dimensions of the VLP particles, extending up to about 20 nm away from the glass-solution interface (**Fig. S1c**), while the contrast to mass conversion was calibrated using small proteins that bind close to the interface. This height change leads to differences in the measured interferometric optical contrast owing to the added optical path difference between the reference light, reflected from the glass-solution interface and the scattered light from the VLP. The measured contrast is therefore a function of the $z$ coordinates through,

$$C \propto 2\frac{|E_{scat}|}{|E_{ref}|}\cos\left(\phi_0 + \Delta OP(z)\right).$$

Where, $|E_{scat}|$ and $|E_{ref}|$ are the real amplitudes of the scattered and reflected fields and $\phi_0$ is an intrinsic phase that includes the Gouy phase shift between the reference field and the field emitted by a point scatterer, as well as any additional phase differences from components of the imaging path such as the partial reflector at the back focal plane and defocusing. Assuming that the microscope was optimised to provide the maximum contrast at the glass-solution interface, we can take $\phi_0(z=0) \approx 0$. $\Delta OP(z)$, is the additional optical path that the scattered field accumulate because of a scatterer that is at distance z above the interface. The optical path difference is given by $\Delta OP = \frac{4\pi\cdot\delta z\cdot n_m}{\lambda}$, and therefore the contribution to the overall contrast as a function of the protein position will be,

$$c_0 \cdot \cos\Delta OP(z) = c_0 \cdot \cos\frac{4\pi\cdot\delta z\cdot n_m}{\lambda}.$$

To estimate the contrast, we calculated the sum $c_{VLP} = c_0\sum_i \cos\frac{4\pi\cdot\delta z_i\cdot n_m}{\lambda}$, where $\delta_{z_i}$, $n_m$, $\lambda$ are the $z$ coordinate of the $i$-th trimeric subunit, the solution refractive index and the wavelength of the laser light source, respectively. Calculation of the expected measured mass, using the above equation as a function of the number of subunits (while increasing the $z$ distances of the trimeric subunits), is shown in **Fig. 1c**. Based on this simple calculation and the measured mass of the mi3 trimer, the expected measured mass of the VLP should be $2015 \pm 24\ k$Da, which agrees well with the measured value. This different between the expected mass and the measured value suggests that the intrinsic phase shift of the setup, $\phi_0 < -0.01\pi$, which indeed close to zero.

***2.4 Solution thermodynamic model (Figure 1d)***

The thermodynamic model for interacting subunits considers two interaction parameters, the monomer-monomer interaction that forms the trimeric subunit, and the trimer-trimer interaction that forms the dodecahedral symmetry. These two interactions are considered by the standard free energy changes $\Delta G_1^\circ$, $\Delta G_2^\circ$, respectively. The law of mass action predicts that the distribution of oligomers will follow,

$$C_n \nu_0 = \sum_i (C_1 \nu_0)^n g_{n,i} \cdot s_{n,i} \cdot \exp\left(-\frac{\Delta G_{n,i}^\circ}{k_B T}\right) \tag{1}$$

Where, $g_{n,i}$ is the number of unique different topological configurations that are composed of $n$ monomeric subunits and includes $c_i$ inter-trimer contacts, $s_{n,i}$ is a kinetic degeneracy factor[4], $\Delta G_{n,i} = 3n \cdot \Delta G_1^\circ + c_i \Delta G_2^\circ$, and $\nu_0$ is the standard state, taken as $1\ M^{-1}$ in our calculation (therefore $C_n$ represent the concentration in M), and $\sum_n nC_n = C_{Total}$, where $C_{Total}$ is the total concentration of mi3-monomers in solution.

The VLP is a regular dodecahedron composed of 20 trimeric subunits, the variation of $g_{n,i}$ with the configuration index, $i$, is of the order of between 1-10 and therefore, $\ln g_{n_i} \ll \Delta G_{1/2}^\circ$ and we can neglect the contribution of this term. This means that the distribution of states can be estimated with high accuracy by considering the most stable state for each oligomeric size and taking only one of its configurations into consideration. This leads to the simplified expression,

$$C_n = (C_1 \nu_0)^n \cdot s_n \cdot \exp\left(-\frac{\Delta G_n^\circ}{k_B T}\right) \tag{2}$$

The law of mass action predicts that at equilibrium and for N>>1, almost all the subunits will be found as either free mi3-subunits or complete VLPs. In our case, this prediction is validated directly by our measured mass histograms (**Fig. S2**). We therefore include three possible states in the thermodynamic model, corresponding to the mi3-monomers, trimers and complete VLPs. We can now write the conservation of mass as,

$$C_1^{Total} \nu_0 = \mathrm{C}_1 \nu_0 + (C_1 \nu_0)^3 \cdot e^{-\frac{3\Delta G_1^\circ + \ln S_{Trimer}}{k_{\mathrm{B}} T}} + (C_1 \nu_0)^{60} \cdot e^{-\frac{20\cdot(3\cdot\Delta G_1{}^\circ + \ln S_{Trimer}) + 30\cdot(\Delta G_2^\circ + \ln S_{VLP})}{k_{\mathrm{B}} T}} \tag{3}$$

where, the degeneracy factors for the trimer and the complete VLPs are $S_{Trimer} = \frac{2^3}{3}$ and $S_{VLP} = \frac{3^{19}}{20}$. The conservation of mass allows us to calculate the concentrations of the three molecular species for each combination of $C_1^{Total}$ and the two interactions free energies.

To extract the standard free energy changes, $\Delta G_{1/2}^\circ$, we fitted the titration at different total protein concentrations (**Fig. 1**). At each iteration, the model assumed the value of the two standard free energies and then solve the relative molar fractions of the different species according to the conservation law, until good agreement between the model and the data has been reached.

#### Supplementary note 3: Dynamic MP measurements on supported lipid bilayers

***3.1 Supported lipid bilayer preparation***

Supported lipid bilayers were prepared using a similar procedure as previously reported[5] with small modifications. In short, phospholipid stocks in chloroform were mixed to form a 5 mM stock solution with a molar composition of 0.05 mM DGS-NTA, 0.1 mM 18:1 PEG550 and 4.85 mM POPC. The stock solution was stored at -20 °C. Before use, 50 µL of the lipid stock solution was added to 200 µL of chloroform in a clean glass tube. The chloroform was evaporated by manually rotating the tube while applying a weak flow of nitrogen, followed by 1 h evaporation under vacuum. Lipids were hydrated by adding 0.5 mL of buffer (20 mM HEPES pH 7.4, 150 mM KCl), followed by 2 cycles of 20 min incubation in a 40 °C water bath, mixing between each cycle. The sealed tube was left at ambient room temperature for at least 2 hours or overnight. The hydrated lipids were tip-sonicated in a 1.5 mL Eppendorf tube and using a 2 mm tip probe at 30% power and 1 sec pulse duration separated by 3 sec waiting time for a total of 10 min sonication time (Vibra-Cell, Sonics & Materials). During sonication, the tube was kept in ice water. The sonicated lipids were centrifuged at 21,130 g for 30 min at 4 °C, before taking 0.4 mL of the supernatant. Cleaned coverslips were treated with oxygen plasma for 3 minutes at 40% power and 0.6 mbar oxygen pressure (Zepto plasma cleaner, Diener Electronic). Immediately following plasma cleaning, a silicon gasket (GBL103280, Grace Bio-Labs) was placed at the center of the coverslip and 30 $\mu$L of buffer (20 mM Tris pH 7.8, 150 mM NaCl, 2 mM $MgCl_2$) followed by 20 $\mu$L of lipids were added and thoroughly mixed in the gasket.

##### *3.2 Preparation of the histidine tag mi3-VLPs for measurements on SLB*

Spytag-polyhistidine peptide (Spy-hist) at a concentration of 270 $\mu M$ in DPBS was mixed with 84 $\mu M$ solution of mi3-VLP (total monomer concentration) at a volume ratio of 2:1, resulting in a large excess of the Spy-hist peptide (180 $\mu M$ vs. 27 $\mu M$) and the mixture was incubated on ice for 3 h. Following incubation, the solution was filtered through a 4 mL, 100 kDa MWCO centrifugal filter (Amicon) at 4000g 8 times to remove the excess peptide. For each round of centrifugation, the 4 mL initial solution was concentrated to 0.1 mL. This resulted in an estimated dilution factor for the excess of Spy-hist peptide of $10^9$. To tether the subunits to the supported lipid bilayers, the tagged VLPs were disassembled by diluting 1 $\mu L$ of the tagged VLP solution into 50 $\mu L$ of 2.5 M of guanidinium thiocyanate. Following about an hour, the disassembled VLPs were rapidly diluted 100-fold into 20 mM HEPES pH 7.4 138 mM NaCl (assembly buffer). The final concentration of tagged mi3-monomers is estimated to be 5 nM, which is much lower than the critical concentration for the formation of the VLP. MP measurements validated the existence of only mi3-monomers and trimers in solution, prior to addition as a solution on top of the supported lipid bilayers.

#### Supplementary note 4: Dynamic MP acquisition and data analysis (Figures 2-3)

##### *4.1 Data acquisition*

Dynamic MP measurements of the two-dimensional assembly reactions of the pentagonal face were performed on a commercial mass photometer (oneMP, Refeyn Ltd.). We used the “medium” field of view ($6.3 \times 9.9$ $\mu m^2$) at the maximum frame rate of 540 Hz and a metapixel pixel size of 77.35 nm after 4×4 pixel binning. Following frame averaging (2 frames), the effective frame rate was 270 Hz. Following the formation of the SLB (**Supplementary note 3.1**) and washing the excess vesicles from the surface with the assembly buffer, 2 nM of tagged subunits were added to the gasket. Through different incubation times, the density of trimeric mi3-subunits was controlled. When the desired density of particles was obtained the solution was replaced with the assembly buffer for at least 5 times, washing away the soluble subunits and the system was allowed to equilibrate for about 30 min. For characterising the thermodynamics (**Fig. 2**), after equilibration, between 3 to 15 movies, 60 sec long, were acquired to collect enough statistics of particles trajectories, depending on the surface density of particles. Each movie was recorded at a different area of the SLB. Before each acquisition, the microscope stage was adjusted to the optimal focus position. Before the acquisition of each data set, a protein standard was measured to calibrate the contrast to mass conversion using the same acquisition parameters.

##### *4.2 Image analysis*

Movies were analysed using a custom-written python package modified from a previously published version[5]. In short, movie processing is divided into 3 steps: image processing, particle detection and contrast fitting. Image processing: Each frame of the 60 sec movie at 270 Hz was normalized to the total detected photoelectron count. To detect the local reflectivity changes originating from light scattered by the diffusing proteins on the SLBs, we subtracted the constant background of the underlying glass roughness by applying a moving median ratiometric imaging analysis approach[2,5]. We chose a 2.2 sec time window for the moving median, suitable for the expected masses and diffusion coefficients of the tethered proteins. To suppress low spatial frequency intensity modulations, originating from rapid laser scanning of the imaged area, we convoluted each frame with a spatial median kernel of size 15×15 pixels, and divided the ratiometric frame by the result. The results of these image processing operations are images similar to the representative frame in **Fig. 2b**. Particle detection: Individual particles were detected above the intrinsic noise of the SLB by cross correlating each frame with a 13X13 pixels kernel of the experimentally obtained point spread function (ePSF) of individual proteins. The ePSF was calculated by averaging individual PSFs of multiple landing events of monodispersed protein solution. Template detection was applied using *match_template* function from *scikit-image* python package. A cutoff value for template matching was set to 0.4. Only pixels whose value was higher than the cutoff values and that were identified as local maxima within a spatial window of 4x4 pixels were considered as detection events. Contrast fitting: Each candidate pixel then serves as the center of a region of interest (ROI) of size 11×11 pixels and the initial guess for the fitting procedure. The contrast of the detected particle was extracted by fitting the *x,y* coordinated of the center of the experimentally normalized (to 1), interpolated ePSF. The x,y positions were found by minimizing the square difference between the defined ROI around the detected particle and the ePSF shifted to *x,y* position. The minimized function is given by,

$$R^2 = \Sigma_{i,j}\left(c(x,y) \times ePSF(x,y)_{i,j} - ROI_{i,j}\right)^2 \tag{4}$$

Where, $i,j$ are the indices of the *ij*-pixel of the ROI and $c(x,y)$ is a scaling factor of the normalised ePSF that minimizes the $R^2$ value at a given *x,y* position. $c_{min}\ (x_{min}\ , y_{min})$ is the reported measured contrast of the protein/complex. The best fitted contrast at each iteration is given by,

$$c(x,y) = \frac{\Sigma_{i,j} ePSF(x,y)_{i,j} \times ROI_{i,j}}{\Sigma_{i,j} ePSF(x,y)_{i,j}^2}. \tag{5}$$

Generating a trajectory from consecutive localisations

Individual successful and consecutive fitting events across adjacent frames were connected into a single molecular trajectory using the same code published and explained previously [5] using the *trackpy* python package.

##### *4.3 Segmenting trajectories using step detection*

To segment each molecular trajectory to its specifically sampled oligomeric states, separated by 120 kDa, for better mass resolution on the histogram level, characterization of the thermodynamic (**Fig. 2**) and for calculation of the average transitions kinetics (**Fig. 3**), we implemented a step detection algorithm[6]. We combined this implementation with a step size threshold of 50 kDa, which is much lower than the known steps of ~120 kDa due to trimer additions and is slightly higher than the noise introduced by the bilayer interface (~40 kDa s.d at 270 Hz). Specifically, the additional introduced mass threshold was used to avoid the consideration of small mass changes that result from the lateral movement of the particles during frame acquisition, leading to different blurring of the PSF and therefore to small contrast variations. Given the intrinsic bilayer noise level of ~40 kDa at frame rate of 270Hz and our interest here in resolving transitions between known measured masses at raw frame rate, we found this threshold to be suitable, with further confidence with it being able to reproduce simulated data of a known transition rate very accurately (**Fig. S3**). Only trajectories longer that 20 frames (74 ms) were considered for segmentation, where shorter trajectories (<20 frames) were considered without segmentation. The minimum segment was restricted to 3 frames (11 ms).

##### *4.4 Extraction of oligomeric mass and diffusion coefficient*

For the calculation of the diffusion coefficient, all detected trajectories and molecular segments were considered similarly. The diffusion coefficient was calculated as previously reported[5], for trajectories longer than 10 frames (37 ms). For shorter trajectories we did not include a measure of the mobility. The molecular mass of each trajectory or segment was calculated by the median value of the mass trajectory. For a given diffusion coefficient, the assigned mass was corrected as to take into account the motion blur that smears the detected and fitted PSF. This smearing effect lowers the fitted contrast by a few percent, depending on the diffusion coefficient of the protein and the mass. The blur correction was validated both experimentally and with simulations for different masses, diffusion coefficients and acquisition parameters as described previously[2,5]. The masses of molecular trajectories to whom a diffusion coefficient was not assigned, were not corrected.

##### *4.5 Plotting mass histograms and calculating surface molar fractions*

To calculate the surface densities of different oligomeric species, we generated weighted mass histograms from the trajectories data set. To avoid noise detection at lower masses, we considered only trajectories longer than 10 frames (37 ms), short segments of long trajectories were included even if their length was shorter than 10 frames. The contribution of each mass trajectory or segment was weighted by its length and the final histogram was divided by the total number of frames per movie and by the detected area. This results in a mass histogram where the *y*-axis represents the averaged number of detected molecular species per area (or surface density). The histograms (**Fig. 2**) were then fitted to a series of five Gaussian functions for the five oligomeric species, from one trimer to the pentagonal ring. The surface concentration of each oligomer was multiplied by the number of its trimeric subunits and the total surface density of trimers was calculated by the sum of all oligomers. Following normalisation, the molar fraction of trimers in each oligomeric state is given by,

$$\mathrm{X}_n = \frac{n\rho_n}{\sum_n n\rho_n} \quad (6)$$

where $\mathrm{X}_n$ is the molar fraction of trimers in an oligomer of size *n* trimers, and $\rho_n$ is the surface density of this oligomer.

#### Supplementary note 5: Characterising two-dimensional interactions

##### *5.1 Thermodynamic characterisation of 2D thermodynamic model for Figure 2d*

A two-dimensional thermodynamic model was used to characterise the distribution of states on the supported lipid bilayer. Here, we assume that the inter-trimer pairwise interaction (**Fig. S4, top**) is identical for all contacts formed, and that there is no cooperativity between binding sites. This means that global fitting included only one fitting parameter, $\Delta G^{\circ}(2D)$, the free energy change in forming an inter-trimers contact (**Fig. 2**). The model therefore includes the pairwise interaction strength and incorporates the kinetic degeneracy for each transition (**Fig. S4**, bottom). We also emphasise that our ability to track individual trimers and oligomers on the surface allows us to directly measure the number density of subunits instead of assuming it, therefore the total density is a measurable rather than an adjustable parameter. In a similar way to solution (**Supplementary note 2.4**), the law of mass action is given by,

$$\rho_{Total} a_0 = \Sigma_{\mathrm{n}=1}^{5} (\rho_1 a_0)^n \cdot s_n \cdot \exp\left(-\frac{\Delta G_n^{\circ}(2D)}{k_B T}\right) \quad (7)$$

where, $\rho_n$ is the number density of trimeric subunits, $a_0$ is the standard state taken as the cross section of the trimers, $s_n$ is the molecular degeneracy resulting from the multivalency, and $\Delta G_n = (n-1) \cdot \Delta G^{\circ}(2D)$ for $n \in \{2,3,4\}$ and $\Delta G_5 = 5\Delta G^{\circ}(2D)$. The titration presented in **Fig. 3d** was fitted to the above equation, to extract the interaction parameter, $\Delta G^{\circ}(2D)$.

***5.2 Quantifying of the effect of dimensionality reduction***

To quantify to which degree the dimensionality reduction enhances the effective interactions, we consider the phenomenological length scale, useful to characterise this enhancement effect, $h = \frac{K_D^{2D}}{K_D^{3D}}$, where $K_D^{2D}$ and $K_D^{3D}$ represent the dissociation constant in 2D (surface) and 3D (solution). The size of this length scale determines whether oligomers form more favourably in solution (3D) or on the SLB. Given that the standard free energy changes of interactions are written in the scale of the molecular dimensions ($\nu_\circ^{-1}$ for 3D and $a_\circ^{-1}$ for 2D, where $\nu_\circ$ and $a_\circ$ are the specific volume and cross section of the protein) this length scale can be written as $h = \frac{\nu_\circ}{a_\circ} \mathrm{e}^{-\frac{\Delta G^\circ(2D) - \Delta G^\circ(3D)}{k_B T}}$. Therefore, enhancement is expected when $h < \frac{\nu_\circ}{a_\circ} = \frac{4}{3} r$, where the equality assumes a globular protein. Reported estimations of $h$ for preferred dimerization on membranes is on the nanometer scale, however vary between different proteins and experimental or theoretical methods[7-10]. Here, we have the opportunity to directly compare the interactions in 3D and 2D on a single molecule level and provide direct quantification of $h$. Given our quantification of the interactions in solution on the membrane we get $h = \frac{0.74 \pm 0.15\ \mu m^{-2}}{2.5 \pm 0.4\ \mu M} = 0.5 \pm 0.1\ nm\ < \frac{4}{3} r_h = 4.4\ nm$, where $r_h$ is the hydrodynamic radius of the trimer calculated using HullRad v10 software[11].

#### Supplementary note 6 : Measurements of oligomeric dwell times

***6.1 Measurements and mass changes detection***

To quantify the dissociation rate constant of the dimer and trimer of trimers ($k_{off}^{dimer}, k_{off}^{trimer}$) presented in **Fig. 3**, we performed three Dynamic-MP experiments at trimer surface densities of 0.25, 0.34 and 0.72 $\mu m^{-2}$. The experiments were perform as described above (**Supplementary note 4.1**), by adding 2 nM of histidine-tagged subunits on top of the SLB. Following an equilibration time of about 30 min and for each bilayer, we consecutively measured 30 different areas on the SLB, each area of dimensions ~7x10 $\mu m^2$, for 1 min, and at an effective frame rate of 270 Hz. The detected molecular trajectories were analysed and segmented as described above (**Supplementary note 4.3**). Following segmentation, segments attributed to dimeric and trimeric oligomers were defined as all mass traces whose median mass falls within the experimental range given by the overall mass distribution of the corresponding oligomer (**Fig. 2 and S5**). Dissociation events for dimers or trimers were defined as any mass change during the molecular trajectory where the final mass is lower than the initial mass and that the absolute mass change is larger than 50 kDa. Theoretically, direct analysis of the resultant distribution of dwell times before dissociation will provide information on the interaction dissociation constant. However, this analysis is prone to several statistical and experimental biases, including: early termination of trajectories due to particles leaving the FOV, termination of molecular trajectories owing to identity switching (wrong trajectory linking results from close proximity of particles below the diffraction limit) and mass fluctuations resulting from close proximity of particles that do not interact. We therefore focus in our analysis on the calculation of the average observed transition rate. Here, $< r_{ij} >$ is the average transition rate from an oligomeric state *i* to any oligomeric state $j$, where $m_j < m_i$, and $m$ the measured mass. Taking the inverse of this rate, $\tau_{ij} =< r_{ij} >^{-1}$, represent the average characteristic time scale for disassembly of oligomer, $i$, or the average dwell time before disassembly occurred. The calculation of the average dissociation rate for the *i*-th oligomer, $< r_i >$ follows,

$$< r_i > = \frac{\sum_{j<i} N_{ij}}{\sum_k t_{i,k}} = \frac{N_{diss.}^{(i)}}{T_{total}^{(i)}}. \quad (8)$$

Here, $N_{ij}$ is the number of detected transitions between state *i* to state *j*, where $j < i$, and $t_{i,k}$ is the total observation time of the *k*-th segment of state *i*. Therefore, the average is given by the total number of disassembly events, $N_{diss.}^{(i)}$ , divided by the total observation time, $T_{total}^{(i)}$. An example of the calculation for representative trace is shown in **Fig. S3**. The average dissociation rate was calculated for the dimeric and trimeric states for each 1-min dynamic-MP movie and converted to the average lifetime, $\tau_i =< r_i >^{-1}$. A distribution of the thirty measured average lifetimes for the two oligomers is shown in **Fig. S5**. The average lifetimes across different surface densities are shown in **Fig. 3** (inset). In addition, since for two dimensional reactions, the kinetic rate constant depends on the local distribution of proteins, the ratio of the average lifetimes of the dimer and trimer was also calculated per movie is shown in **Fig. 3**.

##### *6.2 Dynamic MP simulations for testing the extracted lifetimes*

To validate the procedure to extract the average lifetime, we constructed a simulation framework to generate an approximation for the experimental dynamic MP movies, including known transitions between molecular species.

##### *6.3 Simulation procedure*

*Trajectories generation:* To generate trajectories for a simulation of an experimental 7x10 $\mu m^2$ FOV, a larger FOV of 21x17 $\mu m^2$ with periodic boundary conditions was generated. This way we could keep a constant averaged surface density for individual species, while allowing particle numbers in the field of view to fluctuate. For a given total mi3-trimer surface density, an equilibrium distribution of oligomers was generated using the thermodynamic model described by **equation S7.** From this distribution N random samples of oligomers were taken, where $N/A$ gives the required surface density, with *A*, the area of the FOV. For each oligomer, a 2D diffusion trajectory of length 2 min and temporal resolution of 5.4 kHz (x10 of the measured frame rate) particle movement was generated according to,

$$x_{t_i} = x_{t_{i-1}} + \sqrt{2D\Delta t} \times randn(0,1) \tag{9}$$

$$y_{t_i} = y_{t_{i-1}} + \sqrt{2D\Delta t} \times randn(0,1)$$

Here, $x_{t_i}$ and $y_{t_i}$are the instantaneous position of the particle, $D$ is the diffusion coefficient, $\Delta t$ was set to 0.185 ms (for 5.4 kHz), and randn(0,1) is a randomly drawn number from a standard normal distribution with mean 0 and variance of 1. The initial positions $(x_{t=0,i}, y_{t=0,i})$ were chosen randomly withing the FOV. The diffusion coefficient and the mass of the particles were set according to the oligomer identity and the measured values. This way we can simulate the molecular transitions at similar conditions as the experimental data. To simulate the lifetime of the dimeric oligomer (dimer of trimers), all randomly selected particles that belong to either the mi3-trimers or dimer of trimers where considered together, and the dynamic transitions between mi3-trimers and dimer of trimers were simulated by a given $k_{off}$ that corresponds to the simulated lifetime, and $k_{on} = \frac{k_{off}}{K_D}$, where $K_D$ is the measured dissociation constant for the dimer of trimers (**Fig. 2**). This way, we can simulate the dynamic transitions between the two molecular species while maintaining the overall equilibrated distribution.

For the trajectories classified as either mi3-trimer or dimer of trimers, a transition involved a change of mass from dimer of trimers to mi3-trimer was randomly sampled according to an exponential distribution,

$$\delta t_{dissociation} = \frac{1}{k_{off}} \log{(1/r)} \tag{10}$$

Where, $\delta t_{dissociation}$ is the time difference since the particle first assigned the mass of a dimer of trimers and the transition time, r is a random number drawn from a uniform distribution in the range 0 and 1.

In the same way, stochastic mass transition from a trimer to a dimer of trimers was sampled as,

$$\delta t_{association} = \frac{1}{k_{on}} \log{(1/r)}. \tag{11}$$

At the centre of the large 21x17 $\mu m^2$ FOV, we defined a smaller FOV that is 10% larger than the experimental FOV. Only segments of the trajectories that were occupying this area during the 1 or 2 min of the simulations were saved for further steps.

*Movie generation:* To generate a complete simulated dynamic-MP movie, a 2 min experimental movie of a clean bilayer without added proteins was recorded at 540 Hz and under identical FOV as the data. This movie was used as the background for generating the simulated movies. The trajectories where converted to a noise free movie of the simulated diffusing ePSFs. For every frame, the coordinated of each particle $(x_{ti}, y_{ti})$ was convoluted with an experimentally obtained kernel of the point spread function (ePSF, see **Supplementary note 4.2**). The kernel size was 25x25 pixels, which is equivalent for positioning a particle centered on this position. Since the simulation time resolution is 10 higher than the acquired movie (to simulate motion blurring during acquisition), the amplitude of the convoluted ePSF was set to $\frac{c_i}{10}$, where $c_i$ is the assigned contrast of the PSF (negative) corresponding to the oligomeric state of the simulated particle. The generated array of diffusing ePSFs was then binned 10-fold (summing batches of 10 frames together) to down-sample the simulated movie 10 times, back to the experimental frame rate of 540 Hz. The central area of the simulated movie, similar to the experimental movie, was cut, to fit the size of the experimental FOV, and together with the experimental background movie we generated a simulated movie with the experimental noise components according to,

$$Sim(t,x,y) = BG(t,x,y) + \overline{BG(t,x,y)}_{N=500} \times ePSF(t,x,y)$$

where, $BG(t,x,y)$ is the measured movie of the washed SLB, $\overline{BG(t,x,y)}_{N=500}$ is the same movie following a rolling time average of 500 frames, to suppress background noise and $ePSF(t,x,y)$ is the noise free simulated movie of diffusing ePSFs.

##### *6.4 Extracting lifetimes*

We simulated a series of dynamic MP movies, where the lifetimes of the dimers were set to values between 0.3 and 3.5 seconds, with 5 repeats for each condition. Each movie was analysed with the same analysis code as the experimental movies. The extracted lifetimes were compared to the simulated values to validate the analysis method (**Fig. S3**).

##### *6.5 Estimation of the theoretical 2D diffusion limited rate constant*

To estimate the theoretical limit of the time-dependent two-dimensional forward rate constant, $k^{2D}(t)$, we used the previously derived equation[7],

$$k(t) = 4\pi D \left( \frac{1}{\log\left(\frac{4Dt}{\sigma^2}\right) - 2\gamma} - \frac{\gamma}{\left(\log\left(\frac{4Dt}{\sigma^2}\right) - 2\gamma\right)^2} \right) \quad (12)$$

where $\sigma$ and $D$ are the encounter radius and 2D (experimentally measured **Fig. S6**) diffusion coefficient, and $\gamma$ is the Euler's constant. This equation approximates the rate constant at the long-time limit, where $t \gg \frac{\sigma^2}{D}$. In our case $\sigma = 0.0075\ \mu m$, and $D =\ 2.5\ \mu m^2\ s^{-1}$, which gives $t \gg 45\ \mu s$.

#### <u>Supplementary note 7: Dynamic MP measurements (Figure 4)</u>

##### *7.1 Data acquisition*

Dynamic MP measurements of solution bulk assembly kinetics from tethered pentamers (**Fig. 4**) were performed on the same commercial mass photometer (oneMP, Refeyn Ltd.). Here, acquisition used a custom field of view of size 15.4 × 13.2 $\mu m^2$ to allow maximum statistics and a frame rate of 250 Hz for maximum temporal resolution. Each measurement was for the duration of 60 sec. No further averaging was applied. Following the formation of the SLB and washing excess vesicles from the surface, 2 nM of tagged subunits were added to the gasket. The density of trimeric mi3-subunits was controlled by the incubation time. The solution of the tagged mi3 subunits was replaced with the assembly buffer for at least 5 times, washing away the  subunits present in solution, the system was allowed to equilibrate for about 30 min for the formation of pentamers on the surface. The initial mass distribution was measured and the assembly reaction was initiated by adding an equilibrated solution of the re-assembled untagged-mi3 VLPs at total protein concentration of 88, 44, and 29 nM. The added solution of equilibrated VLPs does not contain the histidine tag modification and therefore particles do not bind the SLB (**Fig. S7**) but the surface-assembled pentamers instead. The assembly of surface pentamers into full assembled VLPs was monitored by acquiring consecutive 1 min movies, each at a different area of the SLB and for approximately 40 min per assembly repeat. At each assembly condition, we repeated the experiment three times. For each repeat, a new SLB was formed and the above procedure was followed. Each repeat of the assembly measurements represents a kinetic measurement of approximately 1000 particles (~20-30 particles per measurement times ~30-40 time points).

##### *7.2 Analysing trajectories to extract mass distributions*

Mass histograms of the dynamic MP experiments shown in Figs. 4 and S9-16 were processed in the same way as described in Supplementary note 4.2 with only one additional step. The approximately 4 times larger FOV used here to increase statistics results in small contrast inhomogeneities across the imaged area. To quantitatively correct these small variations (a few percentages), we performed a standard MP experiment (similar to the procedure described in Supplementary note 2.1) of a monodispersed protein calibrant with a known mass. Using this calibrant we constructed a two-dimensional mapping of the relative variation of the measured mass as a function of the *x,y* position of the imaged area (**Fig. S**8). Following the fitting stage of our dynamic MP analysis procedure (Supplementary note 4.2) we used our relative contrast variation map to correct the measured contrast using the fitted *x,y* position of the measured complex. Since the correction mass is a constant function related to the microscope, that same correction was used for all the measurements that used the same size of the FOV. The experimentally calibrated relative contrast variation is shown in Fig. S8.

##### *7.3 Bulk kinetics modelling*

Aiming at connecting our single complex dynamics data (**Fig. 5**) with the bulk distribution kinetics (**Fig. 4**), we divided the bulk mass histograms into three molecular states: pentamers, complete/close to complete VLP (> 1750 kDa) and intermediates that are between the pentamers and the VLPs in mass (**Fig. S17a-b**). Plotting the fraction of particles in each of these states as a function of time at different solution concentrations of subunits, reveal similar kinetics, however with different timescales (**Fig. S17a**). Here, as expected, following the addition of free trimeric subunits to the solution, the molar fraction of pentamers decrease over time, which correlates to an increase in the fraction of the intermediates. The molar fraction of VLP show a sigmoidal curve, where its increase is connected with the decay of the intermediate fraction. This time variation fits a characteristic curve of consecutive reactions. Informed by our single complex dynamics experiments, showing low probability of disassembly back to the initial pentameric state, in modelling the kinetic rates we included only forward reactions. Fitting this simple model that takes into account the two reactions rates ($r_1$ and $r2$, **Fig. 17b**), results in good agreement with the data (**Fig. S17a**). We attribute the small deviations between the model and the data of the pentamers at the two lowest concentrations to the fact that the concentration of subunits in the solution above the membrane, although assumed constant, is slightly decaying over a period of a few tens of minutes at such low concentrations. Since the perturbation out of equilibrium is small to begin with and the concentrations in solution are very low (17-5.6 nM), small changes in the solution concentrations can result in slowing down or even stopping the conversion from pentamers to intermediates. To quantify concentration dependence, we plotted $r_1$ and $r_2$ across the three conditions on a logarithmic scale (**Fig. S17c**). Linear fits result with slopes of 2.4 $\pm$ 0.1 and 2.1 $\pm$ 0.4 for $r_1$ and $r_2$, respectively. From our single complex dynamics data, we know that the transition between the pentamer to the first stable intermediate state requires addition of three subunits (8-mer), whereas each subsequent face-closing step among intermediates typically requires two subunits. Accordingly, the rates are expected to scale as $r \propto k[Trimer]^n$, where $n \approx 3$ for $r_1$ and $n \approx 2$, which fits very well with our results.

#### Supplementary note 8: Calculation of the grand canonical free energy potential

Computation of the solution grand canonical free energy potential (**Fig. 4** and **S17**) follows previous derivations[4,12-14] and continues the same assumptions introduced in **Eq. S1**. The driving force for the assembly reaction is given by the change in the chemical potential between any oligomer of size *n*, and the trimeric mi3 building block. The molar fraction of the *n*-th oligomer is therefore given by,

$$\frac{\chi_n}{n} = e^{\frac{-n\cdot(\Delta\tilde{\mu}_n{}^{\circ} - \Delta\mu_1)}{k_B T}} \tag{S13}$$

Here, $\Delta\tilde{\mu}_n{}^{\circ}$ is the averaged standard chemical potential change of a trimeric mi3-subunit in an aggregate of size *n*, and $\Delta\mu_1$ is the chemical potential change of soluble mi3-trimer. Since we consider an equilibrated solution added above a passivating bilayer, the concentration of the trimeric mi3 subunits remains constant, and therefore the chemical potential of any oligomer on the surface of the bilayer is calculated relative to $\Delta\mu_1 = \ln\chi_1 \approx Const.$ Assuming a similar set of intermediates to **Eq. S1**, where only the most stable structures for each oligomeric size are considered, this difference can be written as,

$$\Delta\tilde{\mu}_n{}^{\circ} - \Delta\mu_1 = \frac{-c_n\cdot\epsilon - k_B T\cdot ln s_n}{n} - ln\chi_1 \tag{S14}$$

Here, $\epsilon$ is the pairwise interaction free energy gain in units of $k_B T$, $c_n$ is the number of inter-trimers contacts, and $s_n$ is a degeneracy factor[4] similar to that in **Eq. S1**. At the pseudo-critical concentration of $\sim 17\ nM$ of trimers, this difference is shown in **Fig. S17b**. $\Delta\tilde{\mu}_n{}^{\circ}$ is a decaying function of *n*, and the difference in chemical potential with respect to the soluble trimer is always $\geq 0$, as expected at equilibrium where no net driving force for assembly exist. The change in the average coordination number of the incoming trimers as a function of *n* is indicated by the local minima of the curve. This change becomes more apparent when comparing **Eq, S14** without the degeneracy factor (**Fig. S17b** orange curve) to a linear oligomeric growth model (green curve), where addition of each subunit introduces one inter-subunit bond. These local minima, correspond to the topologically closed, stable intermediates.

The relative occurrence of oligomers is better illustrated by examining the grand canonical potential under a fixed chemical potential of subunits (**Fig. S17c**),

$$\Theta_n(\epsilon, \Delta\mu_1) = n\Delta\tilde{\mu}_n{}^{\circ} - n\Delta\mu_1 = -c_n\cdot\epsilon - k_B T\cdot ln s_n - n\cdot ln\chi_1\ . \tag{S15}$$

In solution the chemical potential of 20 trimeric subunits is equal to the chemical potential of the trimers in a complete VLP (**Fig. S17c**, top), and $\Theta_n(\epsilon, \Delta\mu_1)$ represents the potential landscape at chemical equilibrium.

When the solution is brought into contact with the surface pentamers, the grand canonical potential remains unchanged, however, the surface now contains pentamers in excess relative to their equilibrium fraction in solution (disassembly of pentamers is irrelevant as the surface distribution itself is at equilibrium). The relevant chemical driving force is therefore between the pentameric rings on the surface and the complete VLPs, mediated by the constant chemical potential of the trimeric subunits in solution,

$$\Delta\Theta_{5\rightarrow 20}(\epsilon, \Delta\mu_1) = 5\Delta\tilde{\mu}_5{}^{\circ} - 20\Delta\tilde{\mu}_{20}{}^{\circ} + 15\Delta\mu_1 = -25\cdot\epsilon - k_B T\cdot ln\frac{s_5}{s_{20}} + 15\cdot ln\chi_1\ . \tag{S16}$$

Using the fitted value of the free energy per contact (10.7 $k_B T$, **Fig. 1**) and the concentration of trimeric subunits (17, 11.8, 7.4 nM) for the three experimental conditions yields driving forces of approximately 15, 10,3 $k_B T$, respectively (orange marks in **Fig. S17c**).

#### Supplementary note 9: Single-complex assembly experiments

##### *8.1 Coverslip preparation and photolithography*

Confined supported lipid bilayers were prepared using a previously published protocol with modifications[15,16]. Briefly, glass coverslips were cleaned as described in **Supplementary notes 2.2 and 3.1**. Following plasma cleaning, the coverslips were rinsed with MQ water, dried with nitrogen, and fitted with silicone gaskets. The gaskets were filled with 50 µL of 2 µg/mL PLL(20)-g[3.5]-PEG(2) (SuSoS Surface Technologies) and incubated for 30 min at room temperature. After incubation, the coverslips were rinsed with MQ water, dried with nitrogen, and exposed to deep UV light in a mask aligner (Suss MJB4, HgXe 500 W source) for 60 min through a custom-made chrome photolithography mask containing an array of 5 µm-diameter circles. Finally, the coverslips were rinsed with MQ water, dried with nitrogen, and stored at –20 °C for up to three months before use.

##### *8.2 Data acquisition*

Confined supported lipid bilayers were prepared using the photolithographically patterned glass coverslips (**Supplementary note 8.1**) and the supported lipid bilayer preparation procedure (**Supplementary note 3.1**). Formation of the surface pentamers confined to the SLB followed similar protocol as in **Supplementary note 7.1**. For assembly from pentamers to full VLPs, a solution of an equilibrated solution of pre-assembled untagged-mi3VLPs at total protein concentration of 88 or 100 nM was added on top of the confined pentamers. Focus was then found, and the confined supported lipid bilayers were measured for 5 min from the time of solution addition using a OneMP (**Supplementary note 7.1**).

##### *8.3 Data analysis*

Data analysis was done in the same manner as **Supplementary note 4.2**, with additional step for correct long tracking of single VLPs in case where more than one pentagonal ring were confined in the same trap. After automatic trajectory linking, resulting trajectories were further manually linked using frame, position, and contrast values. In a few cases slower mobility of particles close to the edge of the trap affected the ratiometric contrast, due to the median calculation (**Supplementary note 4.2**), we manually found frame periods where the VLPs did not move and reanalysed these with a modified version of ratiometric analysis in which the background is estimated using interpolation of the raw images from 250 frames before immobilisation to 250 frames after immobilisation. The contrast values of the initial pentameric rings were converted to mass by aligning the initial contrast values (first 10s) to the expected pentamer mass, following calibration of the mass to contrast conversion on the same acquisition settings (**Supplementary note 7.1**).

##### *8.4 Mass trajectories analysis*

Dwell times analysis was performed by using the step detection procedure (**Supplementary note 4**) to identify transitions between molecular states, the resultant trajectories are shown in **Fig. S21.** The molecular states were defined according to their expected masses (**Fig. S21**, projected histograms) with a possible error of up to 5% owing to variations in the contrast to mass conversion due to variation in focus position. The trajectories and the molecular transitions were also examined manually to validate the transition times between the stable intermediate structures. In case that a molecular state was not observed in a particular mass trace, its dwell time was set to 0.

#### Supplementary note 10: Stochastic kinetic simulations (Fig. 6b)

VLP assembly dynamics (**Fig. 6b**) were simulated using a simple Gillespie type stochastic kinetic Monte Carlo approach[17]. To model the time evolution of subunit association and dissociation to a growing VLP, we considered the VLP structure as an undirected graph comprising 20 nodes, where each node has a connectivity of 3, corresponding to the interaction sites of the subunits. Therefore, each node represents a trimeric subunit and each edge represents a trimer-trimer interaction. Each simulated trajectory represents the assembly of one VLP particle, under a fixed concentration of trimeric subunits in solution.

Starting with a graph representing the pentagonal ring, in each iteration, all possible potential node addition or removal were enumerated to the current state of the graph. Nodes can be added through the creation of a new edge to the current state of the graph, while removal is only possible by deletion of a single edge that results in an increase in the connected components of the graph. In this case, the connection component that does not contain the original pentameric ring is removed. Physically the later assumption state that subunits that are connected to the growing VLP by more than one interaction, cannot be removed, and therefore topologically closed structures cannot dissociate. The total propensity $a_\circ$ is calculated as the sum of all forward (association) and backward (dissociation) reaction propensities, given by:

$$a_\circ = \Sigma_i^{N_{ass}} a_i^{ass} + \Sigma_i a_i^{diss} \quad (17)$$

where,

$$a_i^{ass} = k_{i,on} [T], \; a_i^{diss} = k_{i,off}. \quad (18)$$

here, $k_{on}$ and $k_{off}$ are the microscopic association and dissociating rate constants per site, and $[T]$ is the trimers molar concentration in solution. Random numbers $r_1, r_2 \in (0,1)$ are sampled from a uniform distribution to determine the next reaction time increment:

$$\tau = \frac{1}{a_\circ} \ln \frac{1}{r_1} \quad (19)$$

and to select the specific event that occurs by the smallest index that satisfy,

$$\Sigma_j a_j > r_2 a_\circ \quad (20)$$

where, $a_j$ is a vector containing the combined propensities for association and dissociation events. According to the chosen event we updated the graph and the recorded time increment. In each trajectory, the simulation proceeds until either the maximum simulation time of 1 h or the completion of the VLP is reached. Simulations were performed at three concentrations 17, 9, and 5.6 nM of trimeric subunits. The association rate and dissociation constants were set to $10^7\ M^{-1}s^{-1}$ and 1 $\mu M$ for considering weak micromolar interactions that are diffusion limited.

#### Supplementary Figures S1-S21

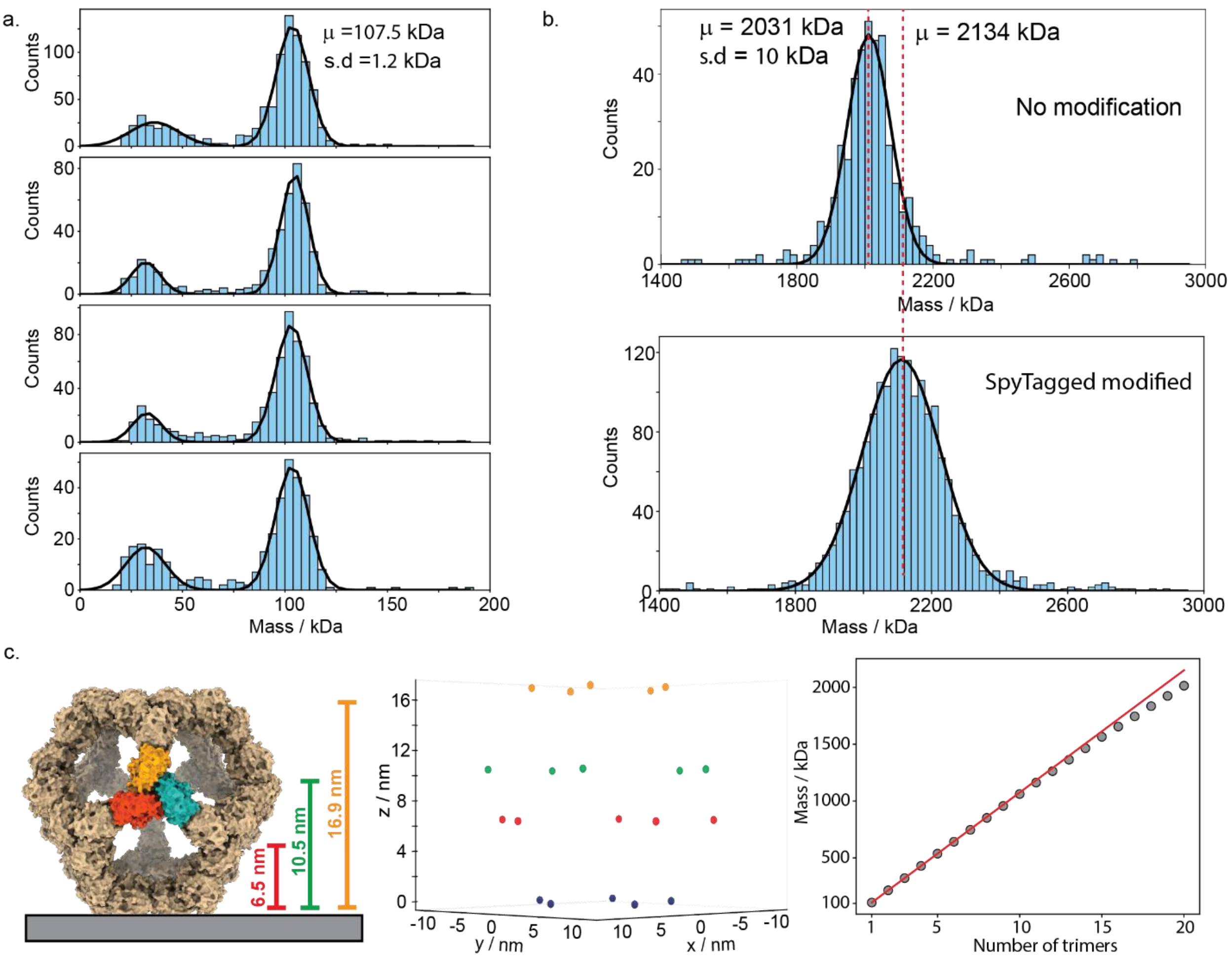

**Figure S1: Mass measurements of mi3-subunits and complete VLPs.** (a) Four representative mass histograms of mi3-monomeric and trimeric subunits following disassembly of the VLP (see Supplementary note 2). (b) Mass histograms of the complete VLPs before disassembly and the complete VLPs following the addition of excess spyTag-polyhistidine peptide (calculated mass of ~3.3 kDa per peptide). The binding of multiple peptides shifts the mass of the VLP by ~100 kDa. (c) The origin of the measured lower mass for the capsid. Left: illustration of the capsid as it expected to land on the glass surface. Owing to the dodecahedral symmetry there are 4 characteristic heights compared to the surface of the glass. The height differences between the subunits and the glass-solution interface are translated to additional phase factors between the reference light and the scattered light, resulting in a lower contrast (grey symbols) than expected from 20 trimers that are bound directly to the glass surface (red curve).

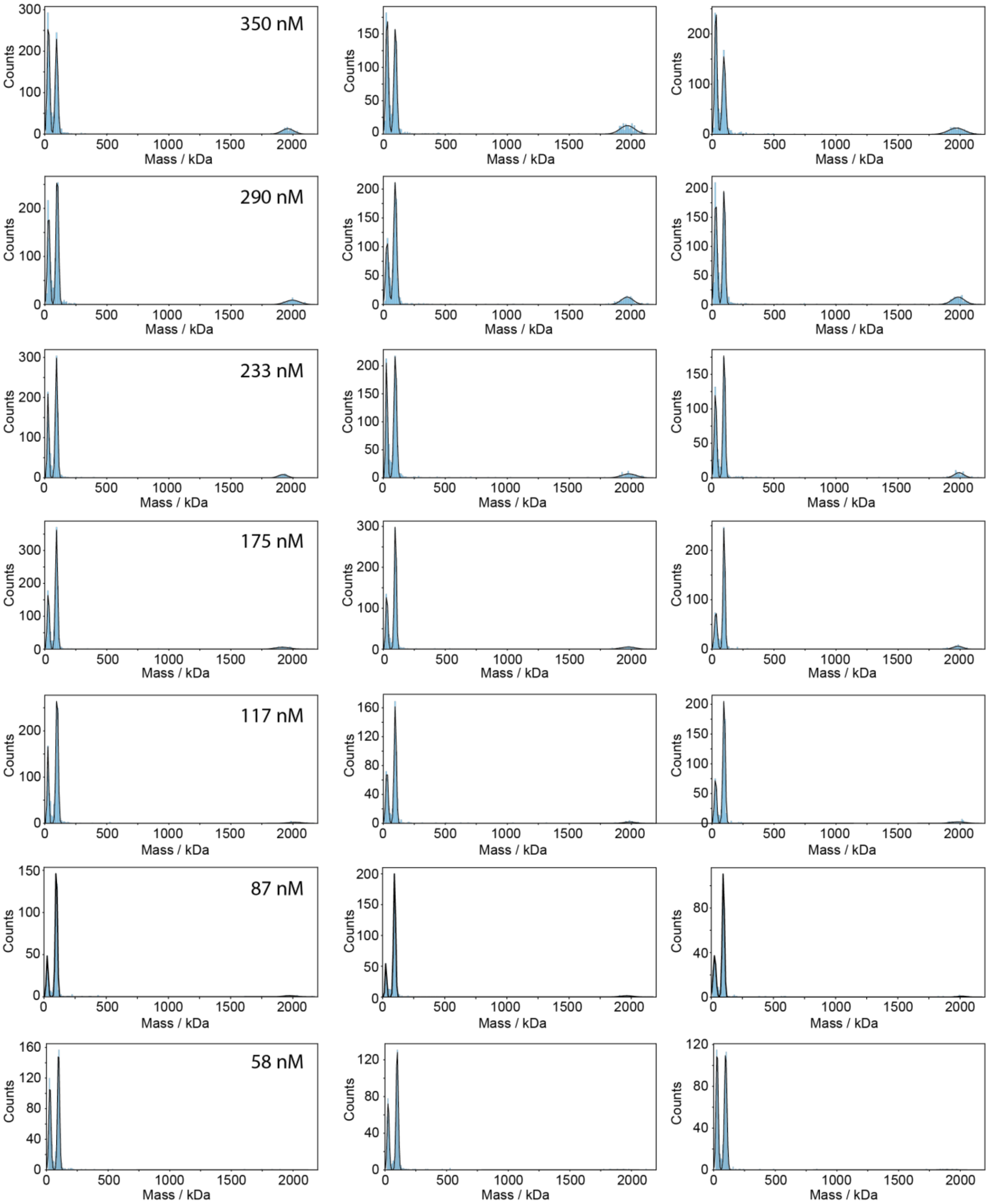

**Figure S2: Measured mass histograms of the assembly titration presented in Figure 1.** Mass histograms of equilibrated subunits in assembly buffer, measured at different total protein concentrations from 58-350 nM (indicated). For each protein concentration (rows), three technical replicates were measured (columns). Black curves correspond to the best fitted three Gaussian functions fitted to each molecular state including the monomer, trimer and complete VLP. The calculated mass fraction presented in **Fig. 1d** represents the average and standard deviation of the three replicates.

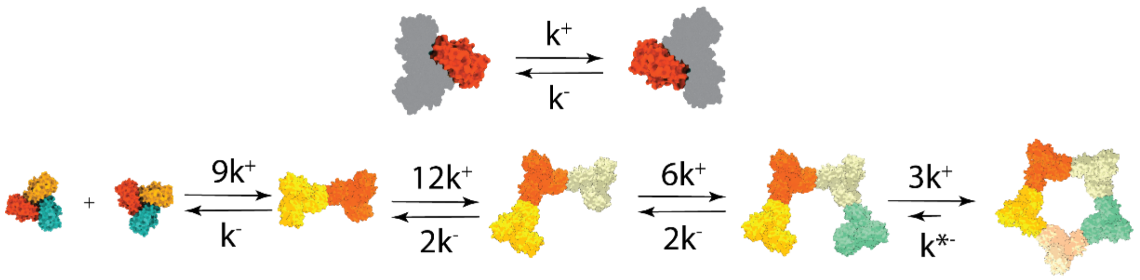

**Figure S3: Schematics of the two-dimensional model illustrating the assembly of the pentagonal ring on SLB surface.** Top: Schematic of the pairwise interaction between one interface of the mi3-trimers, given by $K_D = \frac{k^-}{K^+}$. When considering the interaction between trimers, the kinetic degeneracies should be included, owing to the multivalency. Assuming an isodesmic process, the observed dissociation constant will vary owing to the differences in kinetic factors. The variation in the forward kinetic degeneracy for oligomers larger than 2 is a result of the flat configuration of the growing ring. In this configuration one free interface of the trimer, is oriented to the solution and is not available for 2D interaction. The fundamental interaction free energy change, $\Delta G° = -k_B T \ln \frac{k^-}{\rho \cdot k^+}$ was the only free parameter in the global fit shown in **Fig. 2d.**

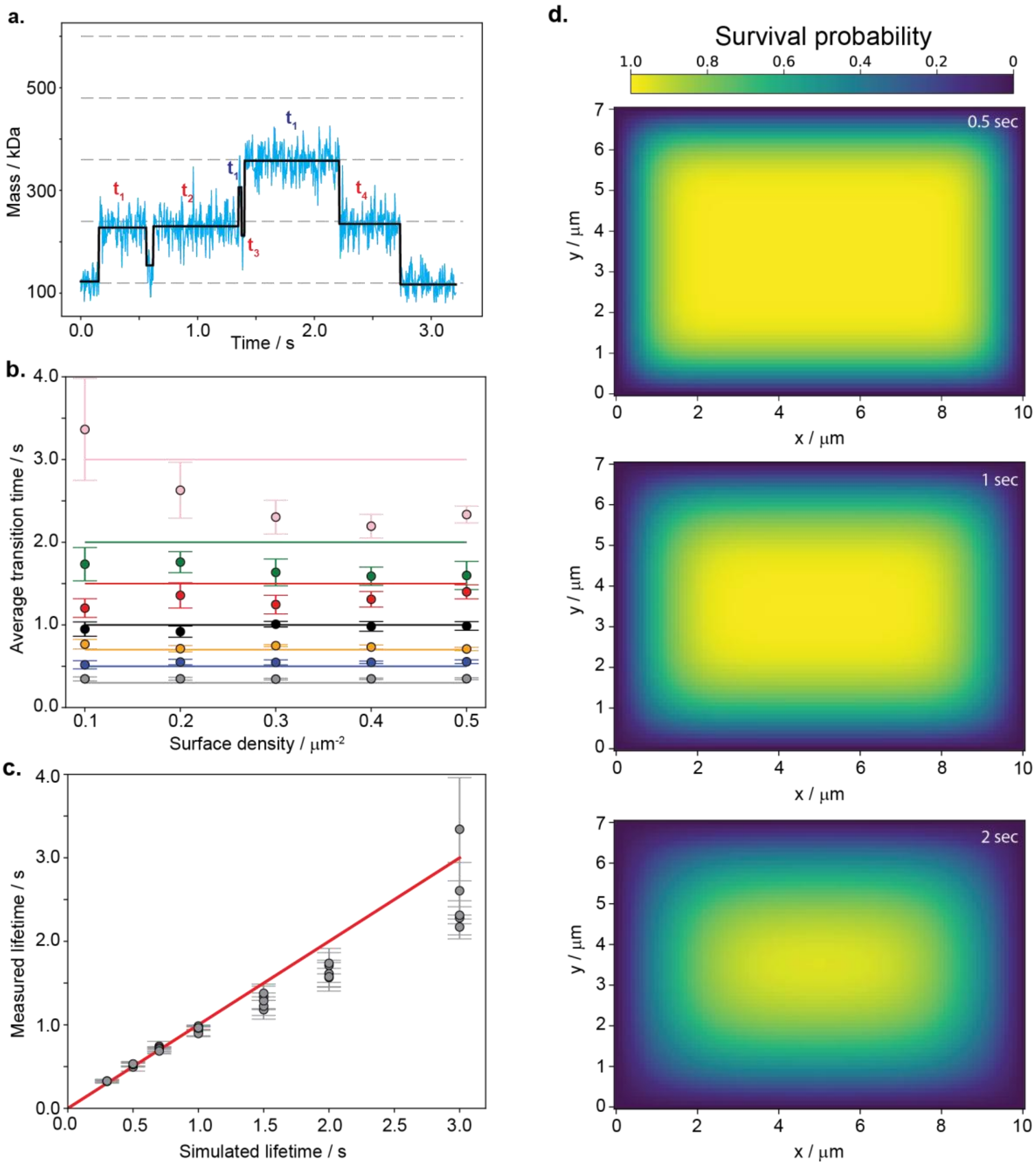

**Figure S4: Step detection and calculation of the average transition times.** (a) Representative example of segmenting a molecular mass trajectory (blue curve) into discrete molecular states (black curve). Red and blue time indices indicate the dwell times for the dime and trimer of mi3-trimers, respectively. Horizontal dashed lines indicate the expected masses of the oligomeric states from mi3-trimer to the pentagonal ring. (b) Average transition times extracted from simulated Dynamic-MP movies (**Supplementary note 6.3**) with defined transition rates between monomer of tmi3-trimers and dimer, across different surface densities. Coloured horizontal lines indicate the simulated transition time, while the scattered symbols and error bars represent the average and standard deviation of values extracted from analysing 5 simulated movies (**Supplementary note 4**). The colours of the scattered symbols correspond to the horizontal line colours. (c) Analysed average lifetime as a function of the simulated lifetimes. Scattered symbols and error bars indicate the average and standard deviation per surface density. (d) Calculation of the probability of a dimer of mi3-trimers to stay in the FOV for the indicated (top right) time period as a function of its initial position. The diffusion coefficient used for calculation was set to the measured value of 0.6 $\mu m^2/s$.

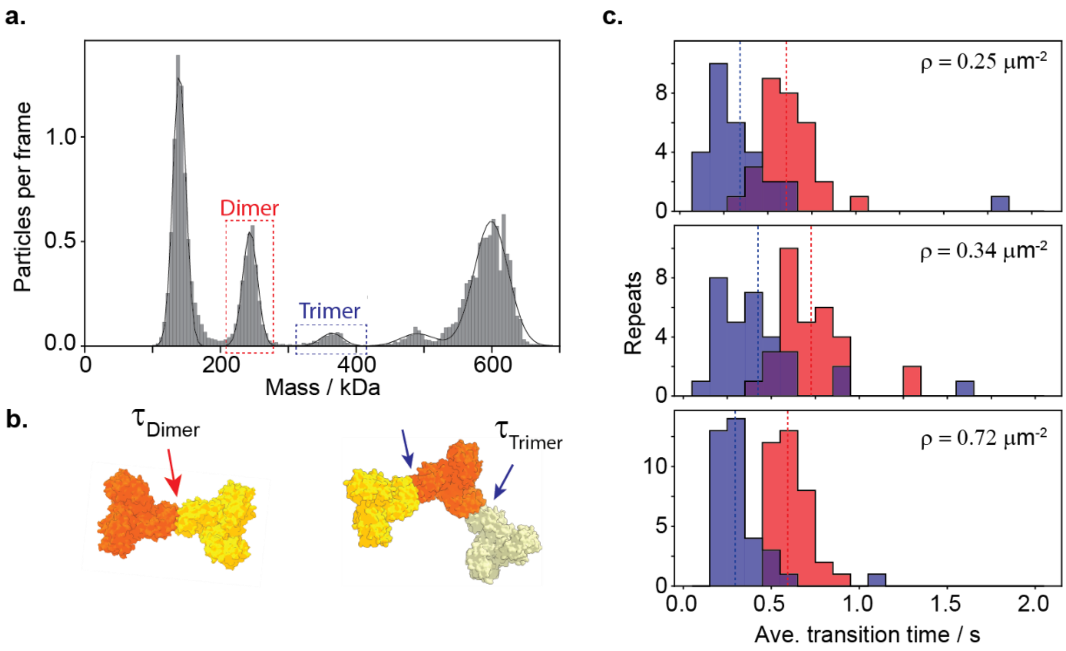

**Figure S5: Quantifying the dwell time of the dimer and trimer on the surface of the SLB.** (a) Representative measured mass histogram (grey bars) of the 5 oligomeric states on the surface of the SLB. Black curve corresponds to the sum of the fitted Gaussian functions for the 5 mass peaks. Red and blue boxes represent the mass ranges that were used to identify dimeric and trimeric oligomers for calculating their averaged dwell times (see **Supplementary note 6**). (b) Illustration showing the origin of the factor if 2 shorter dwell time of the trimer, stemming from the possible dissociation of 2 vs 1 equivalent and independent interacting interfaces. (c) Histograms of the average transition time for disassembly representing 30 independent 1 min dynamic-MP movies. The average surface particle density of each set of experiments is indicated. Red and blue histograms correspond to the dimer and trimers, respectively.

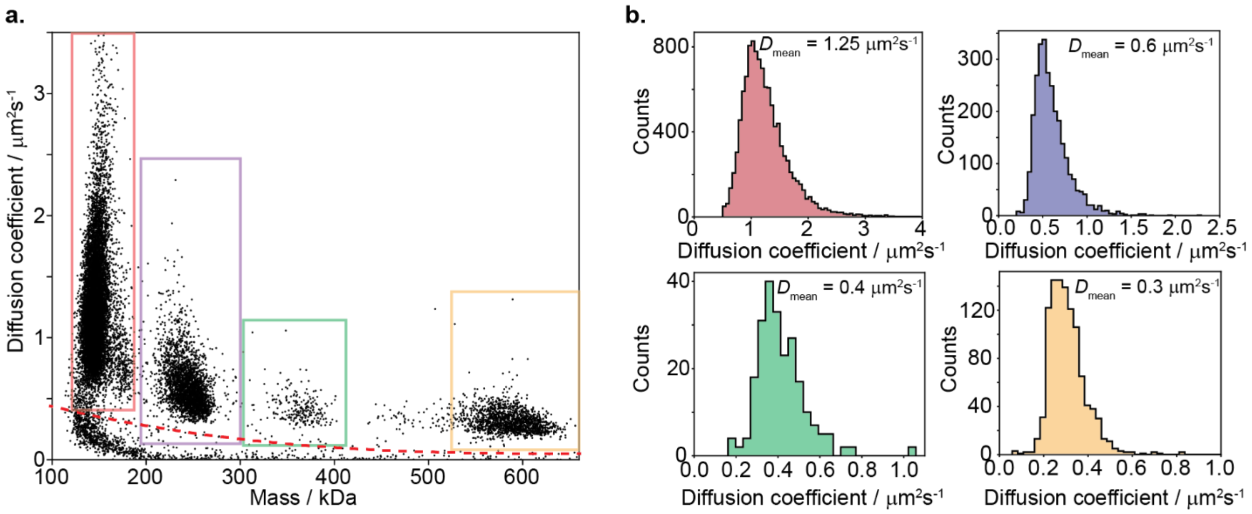

**Figure S6: Mass and diffusion measurement using dynamic-MP**. (a) Representative Mass-Diffusion scatter plot for a 1 min Dynamic-MP movie of the assembly of his-tagged mi3-trimers on an SLB (**Supplementary note 4**). Each point in the plot represents a molecular trajectory segment. Coloured boxes indicate the mass and diffusion ranges corresponds to the mi3-trimer (red), dimer of trimers (purple), trimer of trimers (green) and the pentagonal ring (orange). Red dashed curve corresponds to a threshold below which the trajectories are considered to represent noise from the SLB. (b) Diffusion coefficient histograms, corresponding to the data shown in the coloured boxes in (a). The mean values of the diffusion coefficient distributions are indicated.

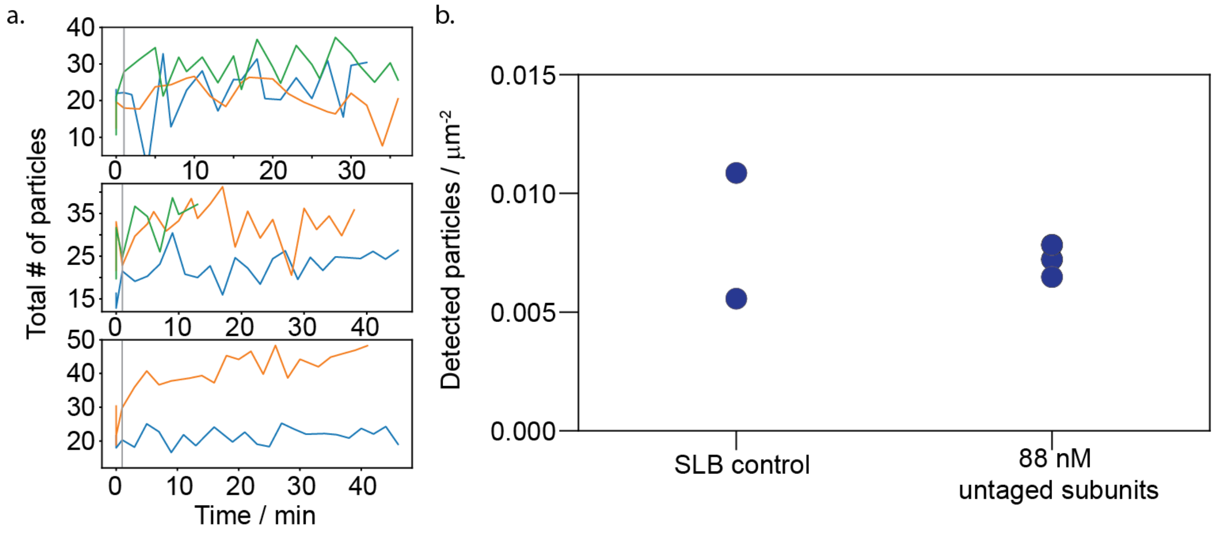

**Figure S7: Characterisation of mi3-subunits non-specific binding to the SLB during dynamic-MP measurements.** (a) The total number of particles observed on the SLB as a function of time during the experiments described in **Supplementary note 7** and presented **in Fig. 4** and **S9-16**. Each colour represents a technical repeat, where the top, middle and bottom panels correspond to the experiments done at total mi3-monomer solution concentrations of 88, 44, and 29 nM. (b) Quantification of nonspecific binding to clean SLBs showing the same and very low number of detected particles before (control) and after the addition of a solution containing a total of 88 nM of mi3-monomers.

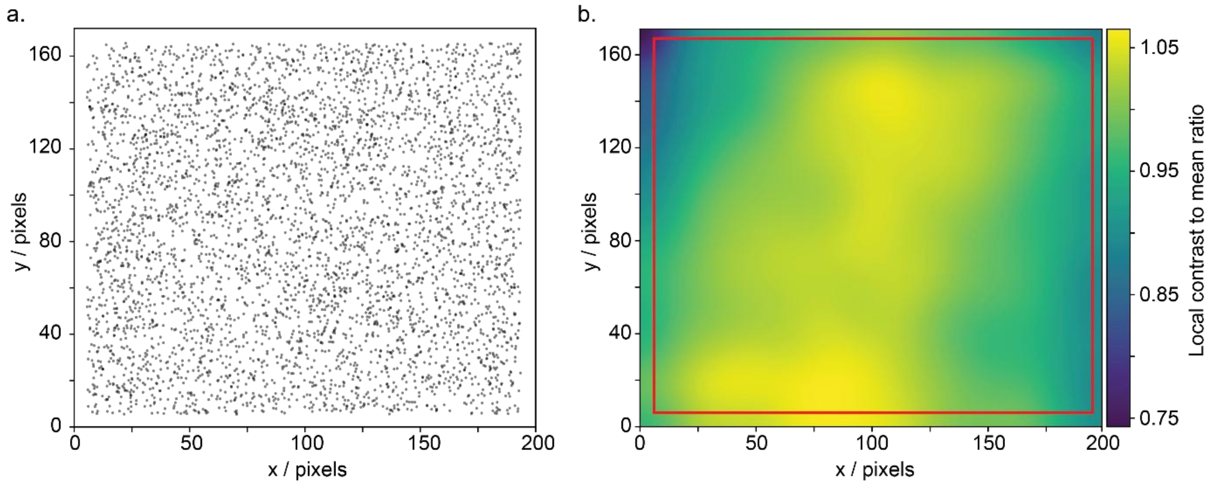

**Figure S8: Spatial contrast correction map for the large FOV used for assembly experiments on continuous BLB.** (a) Binding map of the citrate synthase protein as a function of x-y position. (b) Smoothed and normalised event - contrast map as a function of landing events positions. The red box indicates the area of the FoV utilised for tracking and quantification of the molecular masses of assembling VLPs. The map shows that except for the edges of the FOV, the contrast correction as a function of position is lower than 5%.

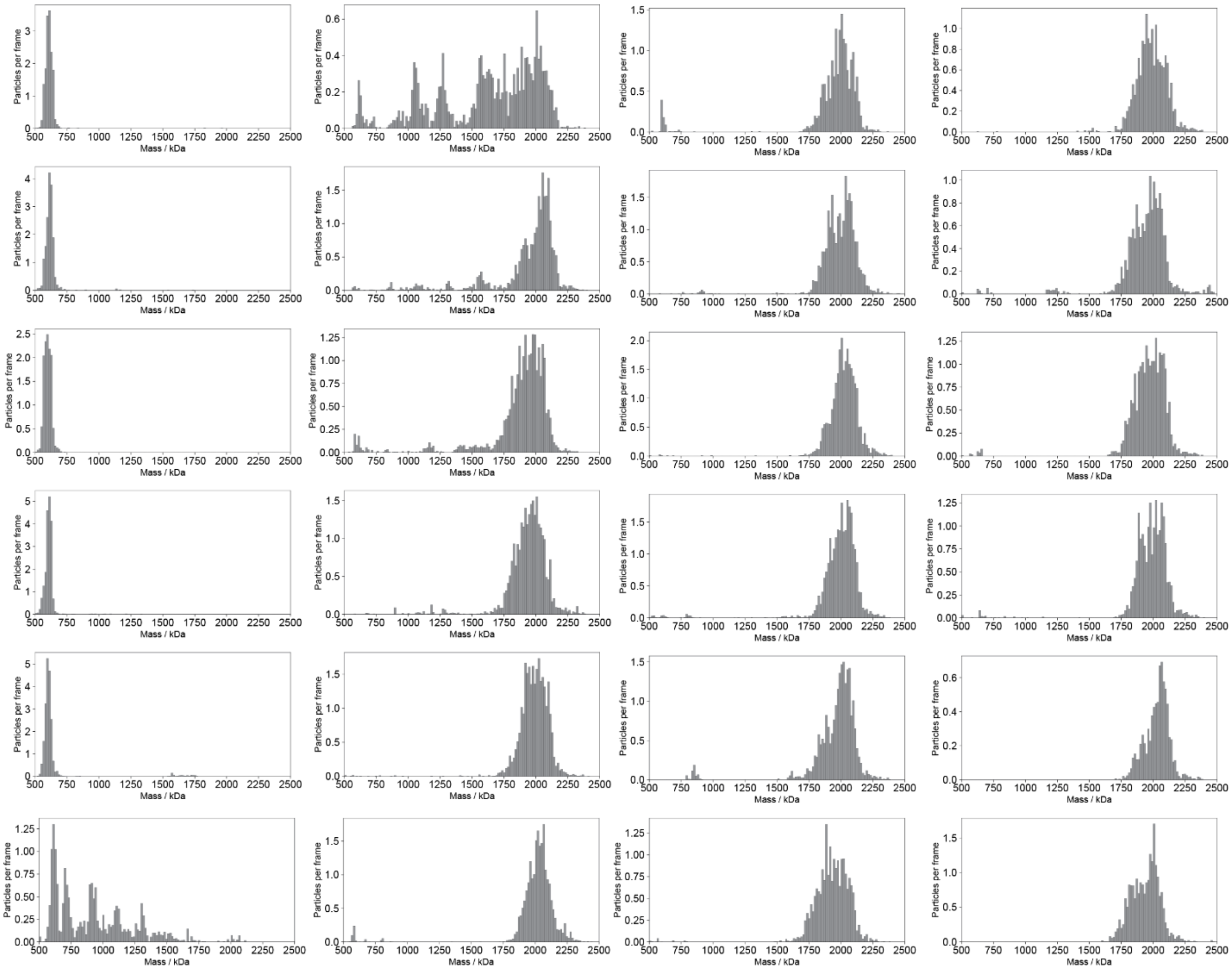

**Figure S9: Bulk assembly mass histograms for assembly reaction at 88 nM of soluble mi3 protein concentration: Repeat 1**. Measured mass histograms for assembly reaction at 88 nM total protein concentrations in solution. Each mass distribution was extracted from a 60 sec Dynamic-MP movie, acquired as described in **Supplementary note 7**. The first 5 histograms represent the reference initial state of the pentagonal ring on the SLB before mi-3 subunits were added to solution.

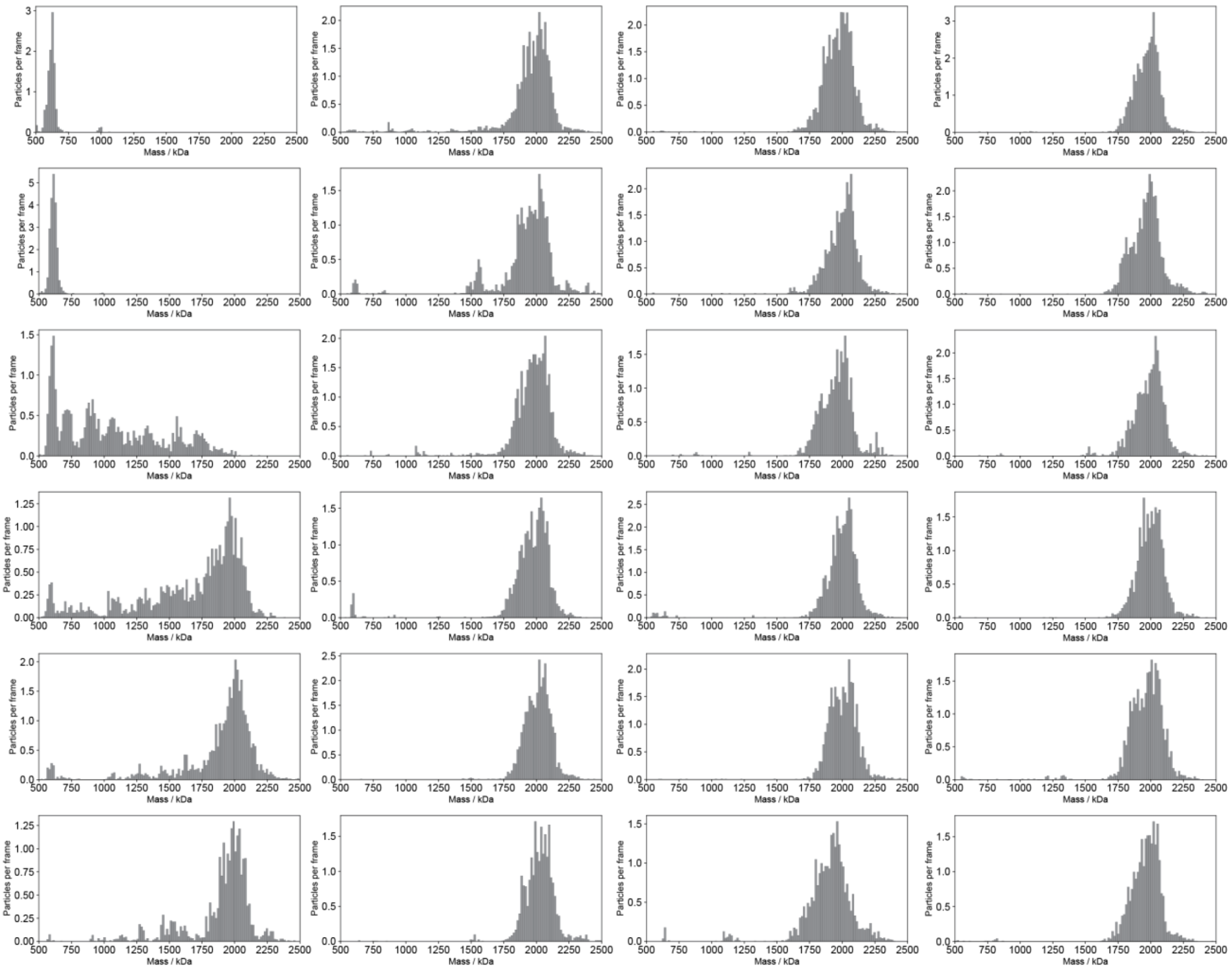

**Figure S10: Bulk assembly mass histograms for assembly reaction at 88 nM of soluble mi3 protein concentration: Repeat 2.**

Measured mass histograms for assembly reaction at 88 nM total protein concentrations in solution. Each mass distribution was extracted from a 60 sec Dynamic-MP movie, acquired as described in **Supplementary note 7**. The first 2 histograms represent the reference initial state of the pentagonal ring on the SLB before mi-3 subunits were added to solution.

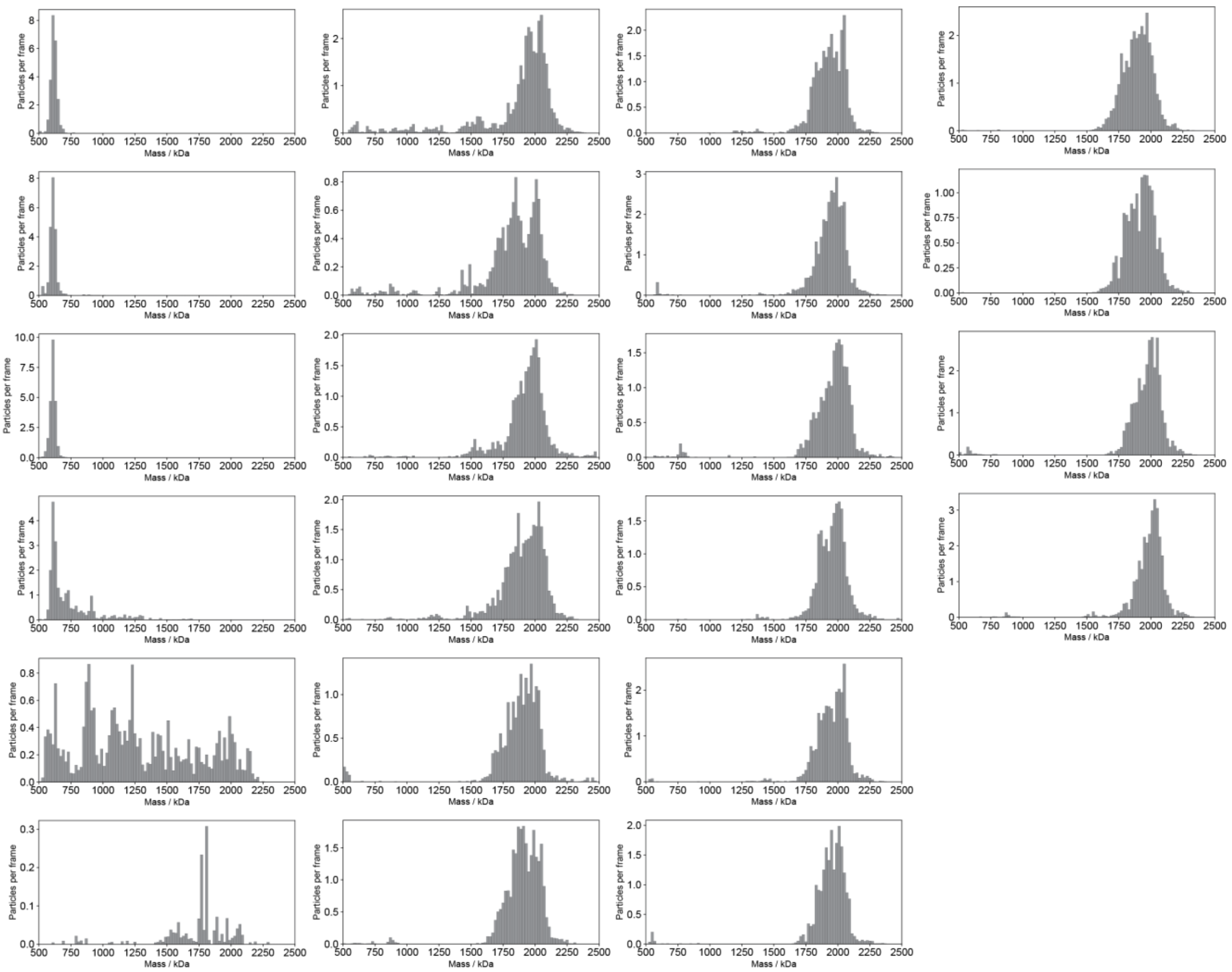

**Figure S11: Bulk assembly mass histograms for assembly reaction at 88 nM of soluble mi3 protein concentration: Repeat 2.**

Measured mass histograms for assembly reaction at 88 nM total protein concentrations in solution. Each mass distribution was extracted from a 60 sec Dynamic-MP movie, acquired as described in **Supplementary note 7**. The first 3 histograms represent the reference initial state of the pentagonal ring on the SLB before mi-3 subunits were added to solution.

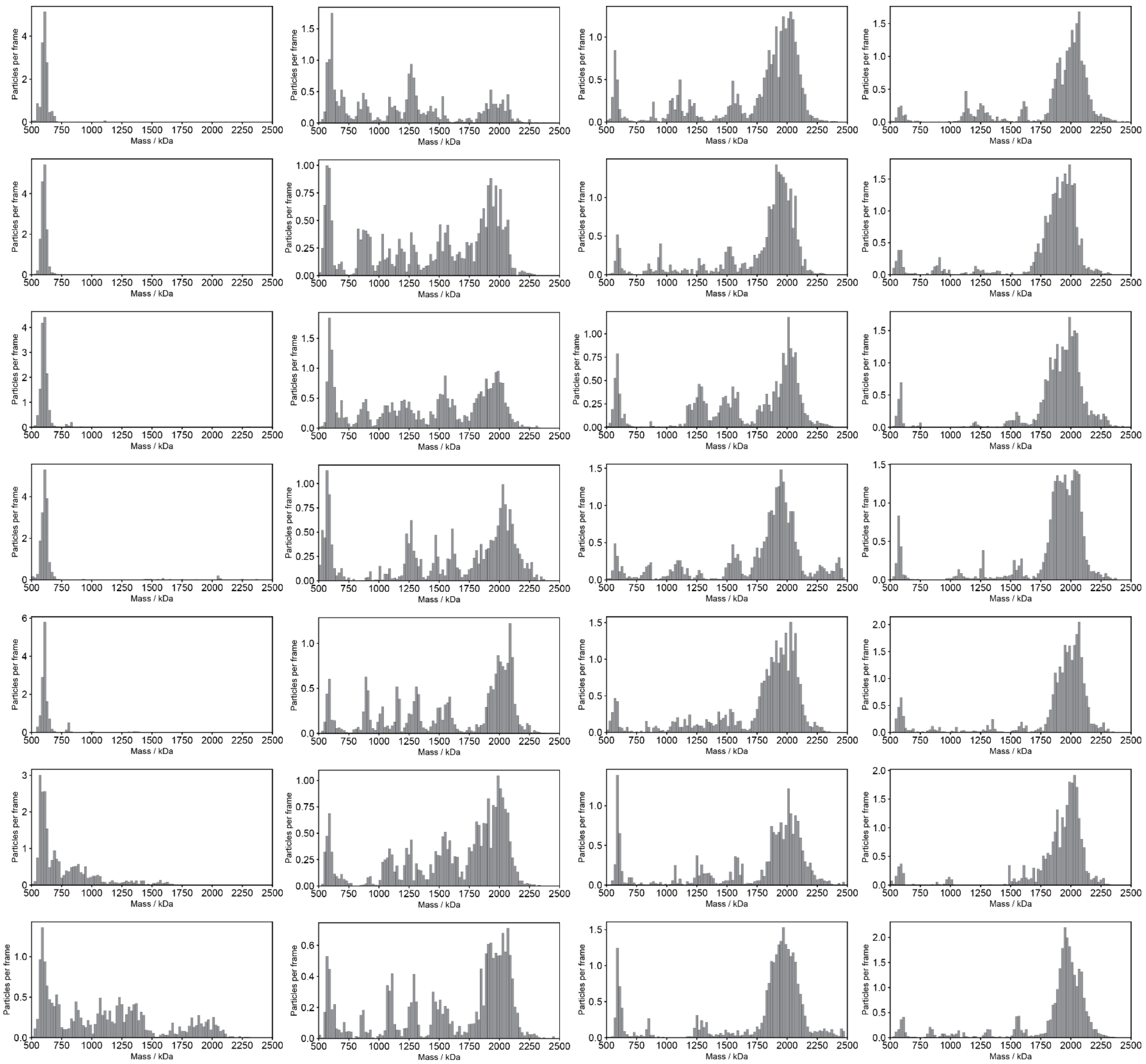

**Figure S12: Bulk assembly mass histograms for assembly reaction at 44 nM of soluble mi3 protein concentration: Repeat 1.**

Measured mass histograms for assembly reaction at 44 nM total protein concentrations in solution. Each mass distribution was extracted from a 60 sec Dynamic-MP movie, acquired as described in **Supplementary note 7**. The first 5 histograms represent the reference initial state of the pentagonal ring on the SLB before mi-3 subunits were added to solution.

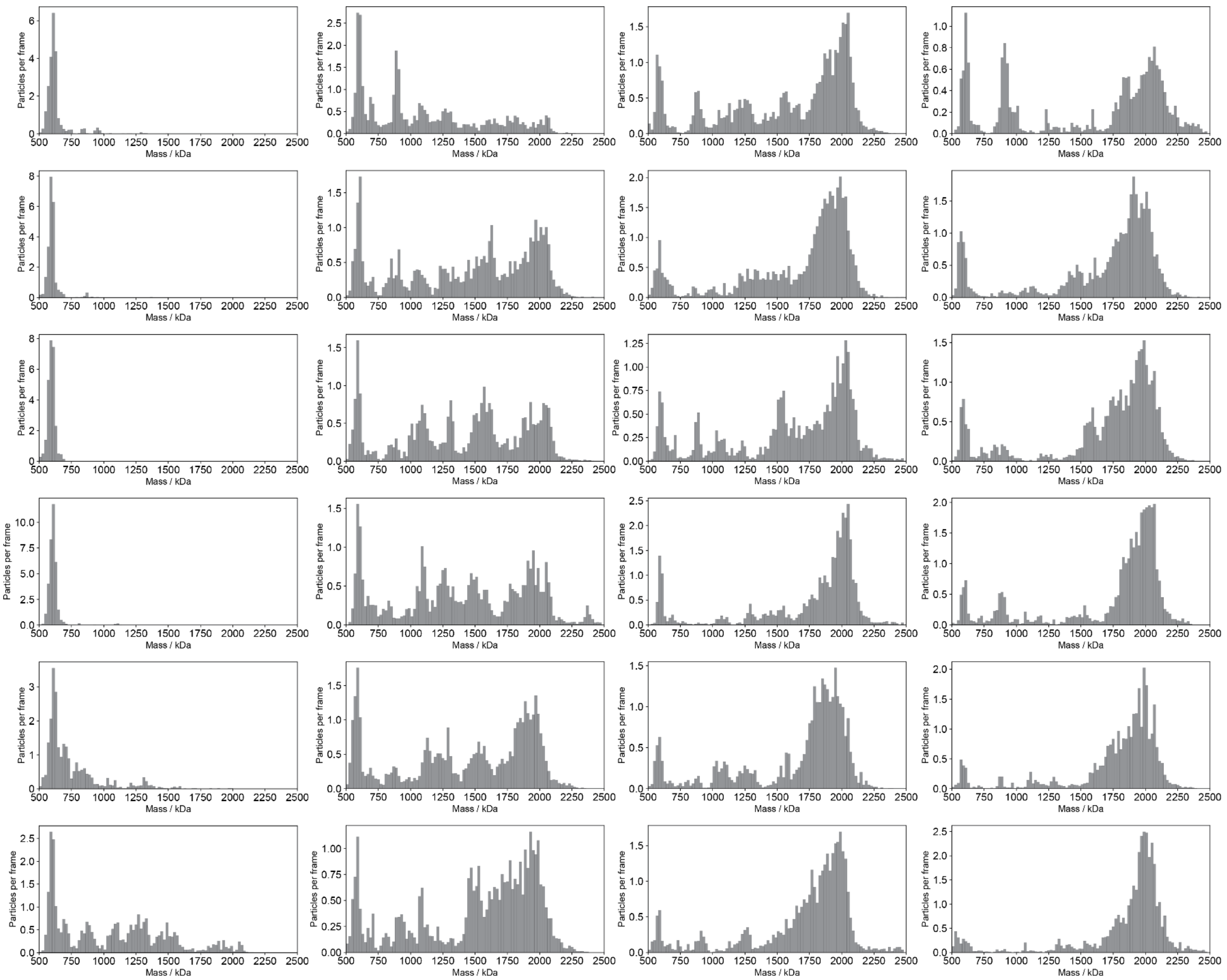

**Figure S13: Bulk assembly mass histograms for assembly reaction at 44 nM of soluble mi3 protein concentration: Repeat 2.**
Measured mass histograms for assembly reaction at 44 nM total protein concentrations in solution. Each mass distribution was extracted from a 60 sec Dynamic-MP movie, acquired as described in **Supplementary note 7**. The first 4 histograms represent the reference initial state of the pentagonal ring on the SLB before mi-3 subunits were added to solution.

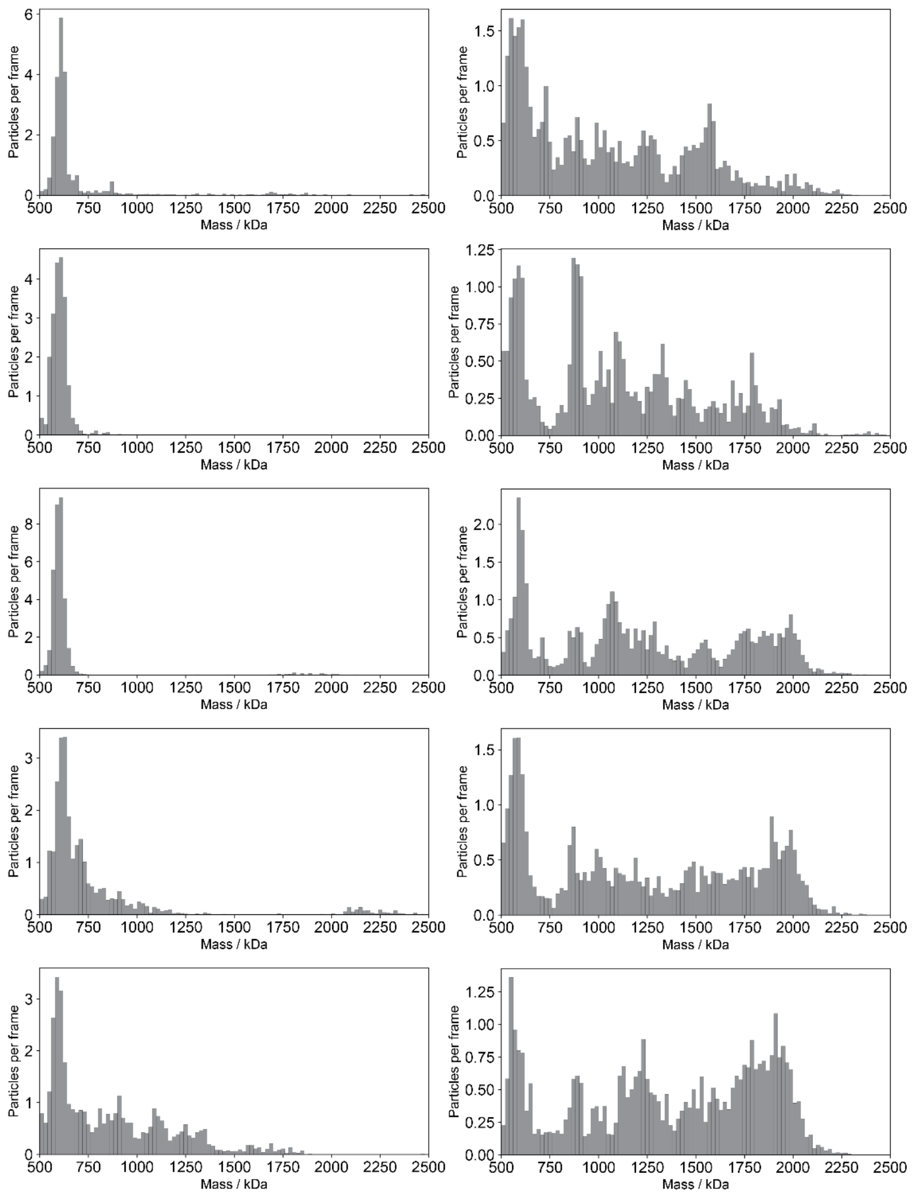

**Figure S14: Bulk assembly mass histograms for assembly reaction at 44 nM of soluble mi3 protein concentration: Repeat 3.**

Measured mass histograms for assembly reaction at 44 nM total protein concentrations in solution. Each mass distribution was extracted from a 60 sec Dynamic-MP movie, acquired as described in **Supplementary note 7**. The first 3 histograms represent the reference initial state of the pentagonal ring on the SLB before mi-3 subunits were added to solution.

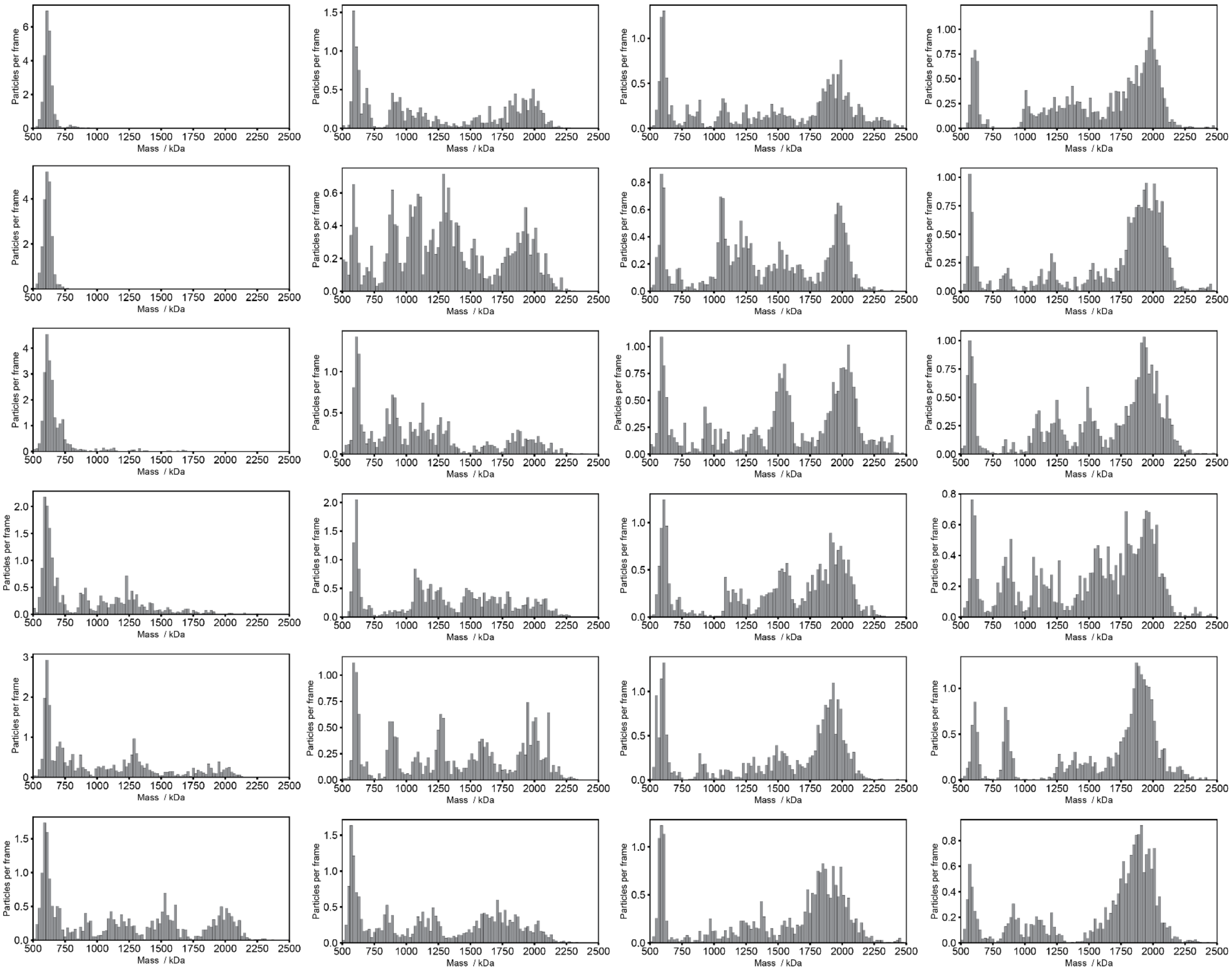

**Figure S15: Bulk assembly mass histograms for an assembly reaction at 29 nM of soluble mi3 protein concentration: Repeat 1.**

Measured mass histograms for assembly reaction at 29 nM total protein concentrations in solution. Each mass distribution was extracted from a 60 sec Dynamic-MP movie, acquired as described in **Supplementary note 7**. The first 2 histograms represent the reference initial state of the pentagonal ring on the SLB before mi-3 subunits were added to solution.

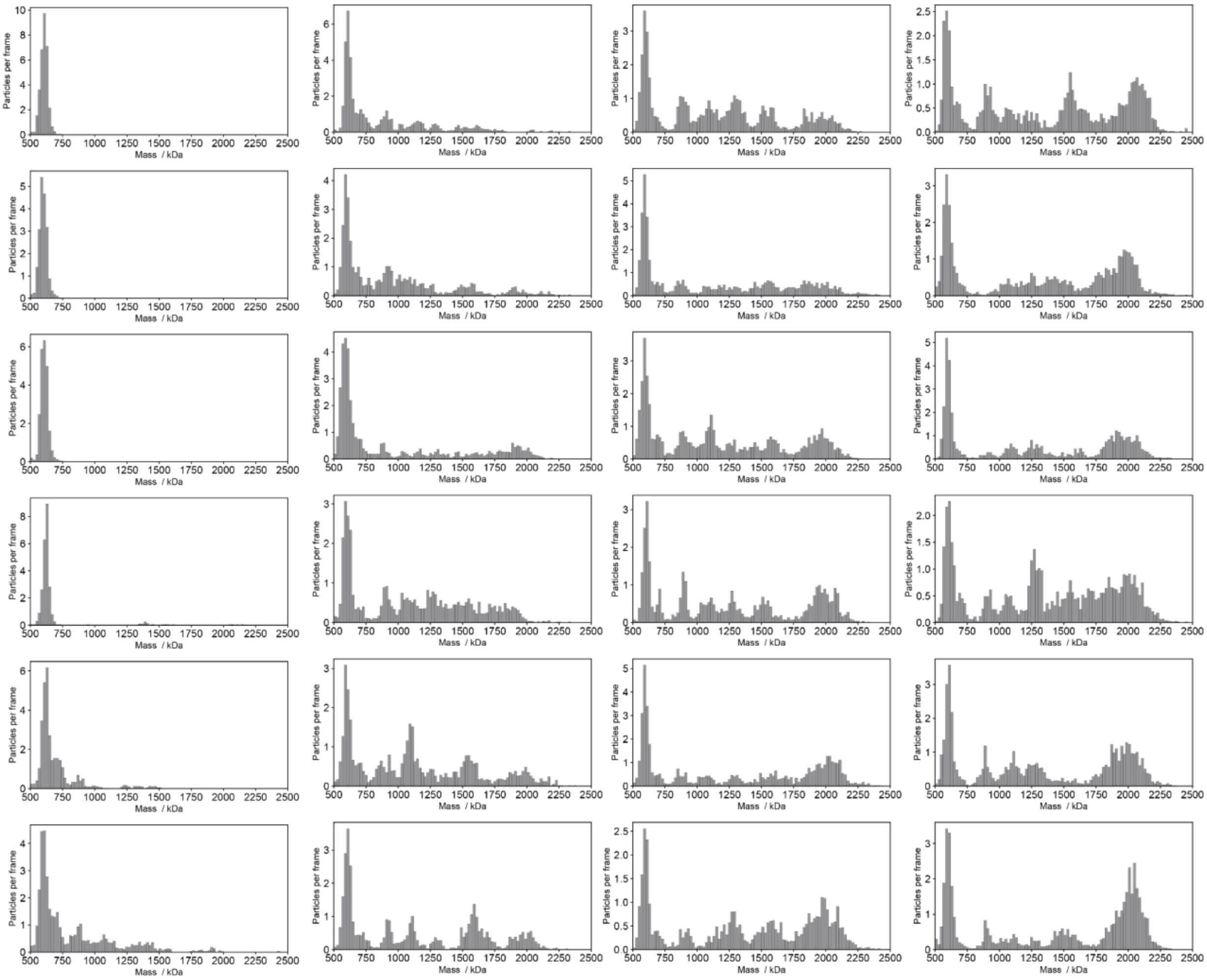

**Figure S16: Bulk assembly mass histograms for an assembly reaction at 29 nM of soluble mi3 protein concentration: Repeat 2.**
Measured mass histograms for assembly reaction at 29 nM total protein concentrations in solution. Each mass distribution was extracted from a 60 sec Dynamic-MP movie, acquired as described in **Supplementary note 7**. The first 4 histograms represent the reference initial state of the pentagonal ring on the SLB before additional mi-3 subunits were added to solution.

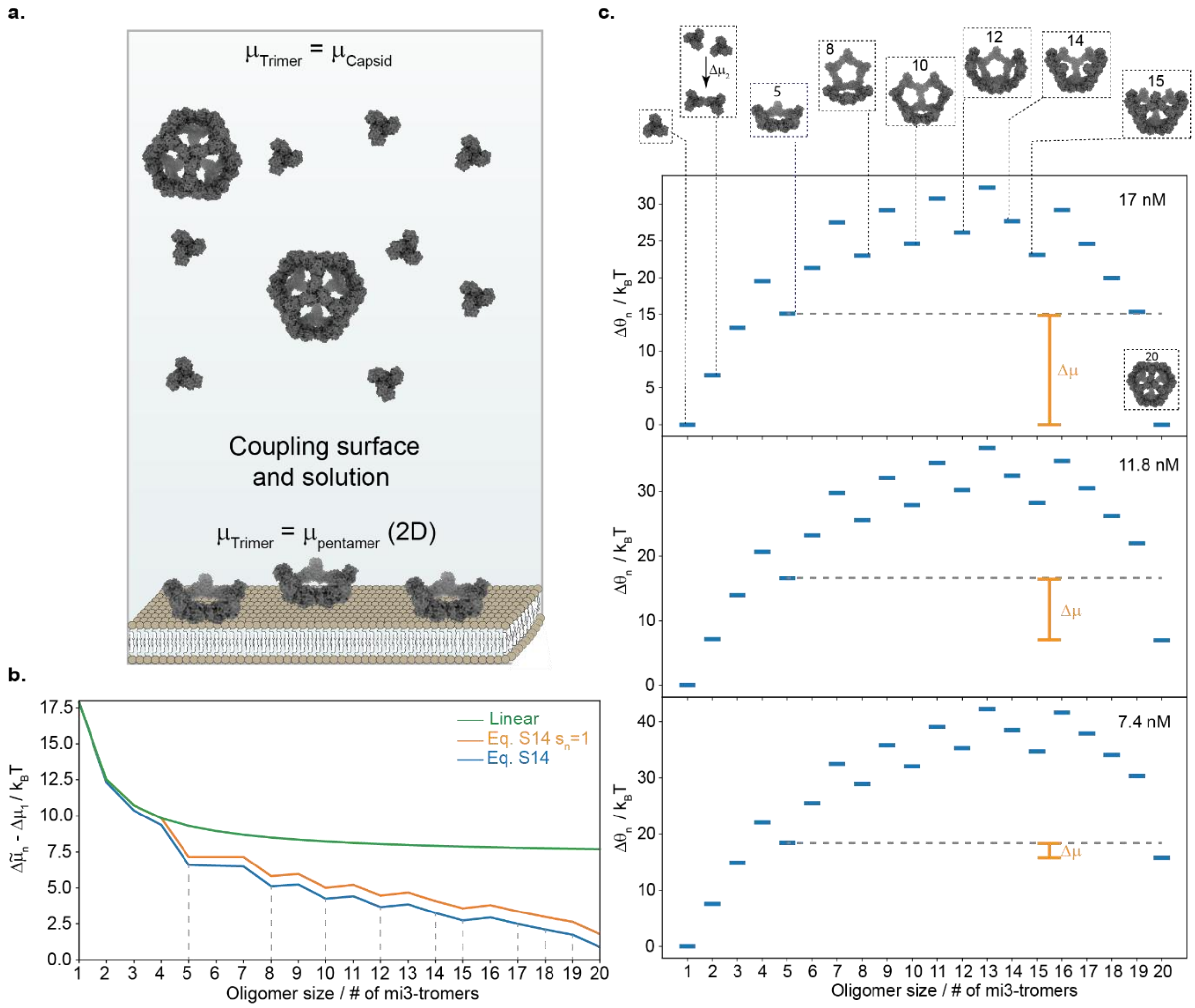

**Figure S17: Calculation of the grand canonical free energy landscape.** (a) Illustration of the experimental procedure used to generate the driving force for the assembly reaction. The subunits on the SLB occupy their equilibrium distributions composed predominantly of pentagonal rings in chemical equilibrium with surface tethered mi3-trimers. Upon addition of the equilibrated solution mixture, composed of soluble mi3-monomers, trimers, and complete VLPs, assembly from the pentameric state becomes energetically favourable, and the pentagonal rings on the surface are elongated through binding of trimeric subunits from solution. (b) The chemical potential difference between a trimer within an oligomer of size *n* and a free trimer in solution, shown as a function of oligomer size for linear oligomers (green), mi3-VLPs oligomers without the degeneracy factor (orange) and mi3-VLPs oligomers with the degeneracy factor (blue) (see **Eq. S14**). (c) The normalised grand canonical free energy potential (**Eq. S15**) at three concentrations of mi3-trimers in solution, corresponding to the experimental mi3-monomer concentrations of 88, 44, and 29 nM. Although there is no driving force for assembly between the trimeric state and the complete VLP, coupling the solution and surface systems and initiating from the pentameric ring, introduces a small driving force (orange). Illustrations depict the molecular structures of the topologically closed configurations.

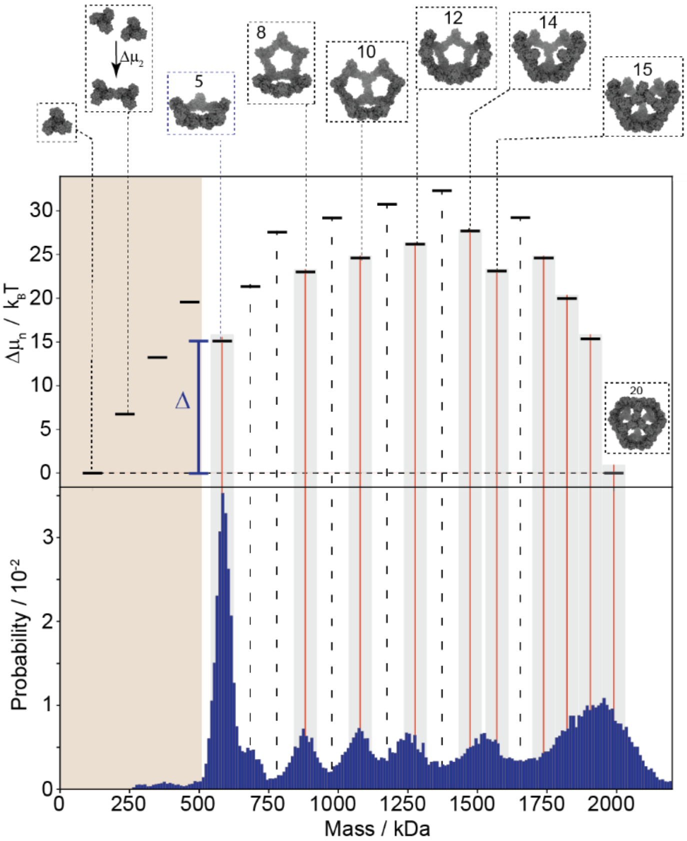

**Figure S18: Comparing the bulk distribution of states for assembly at 29 nM mi3-monomer concentration with the free energy potential.** Top part shows the same minimum grand-canonical free energy landscape shown in **Fig. 4c**. Topologically closed structures that occupy the local free energy minima are shown based on the atomic structure of the VLP (PDB ID: 7b3y), indicated also by solid red vertical lines, while unstable configurations are indicated by black dashed vertical lines. Bottom part corresponds to summing the measured mass histograms (**Figs. S15-16**) at intermediate time points along the assembly reactions (4-30 min) and for 2 technical repeats at 29 nM total protein concentrations in solution.

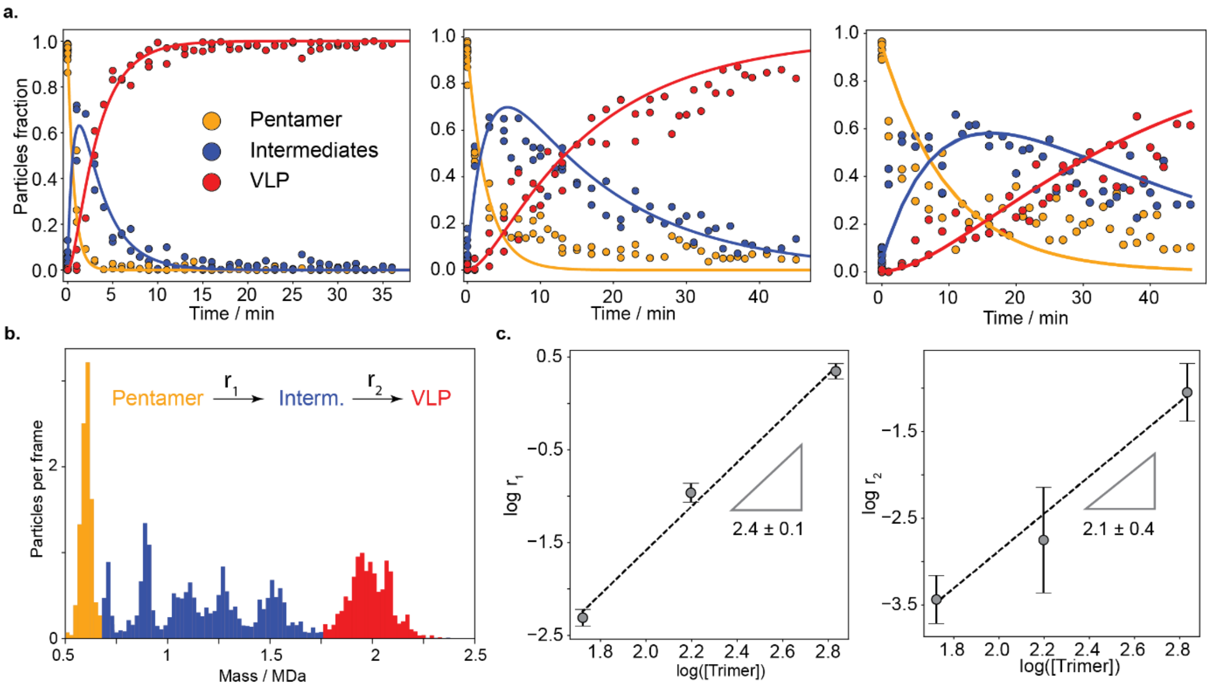

**Figure S19: Clustering the bulk assembly dynamics into three states fits an irreversible kinetic model.** (a) Surface particle fractions as a function of time for the bulk assembly experiments described in **Fig. 4** and **Supplementary note 7**. Normalised particle fractions (symbols) were clustered into three molecular states: the initial pentagonal ring (Pentamers, orange), intermediate structures (Intermediates, blue) and complete/near complete VLPs (VLP, red). The solution total concentrations of mi3-monomers were 88 nM (left), 44 nM (centre) and 29 nM (right). Curves at the corresponding colours indicate the best fitted kinetic model, considering two effective irreversible consecutive reactions, connecting these three states, assuming constant protein concentration in solution. (b) Representative example of the clustering of the Dynamic-MP mass distribution presented in **Fig. 4b and S9-16** into the three molecular states. Legends indicate the expected irreversible kinetic model assuming constant protein concentration in solution and therefore constant effective rates. c. Scaling of the measured reactions rates ($r_{1/2}$) with subunit solution concentration, shown on a semilogarithmic scale, the slope of the curve is indicated. Symbols and error bars correspond to the averaged fitted rates and their standard deviations based on 2-3 replicate for each assembly condition.

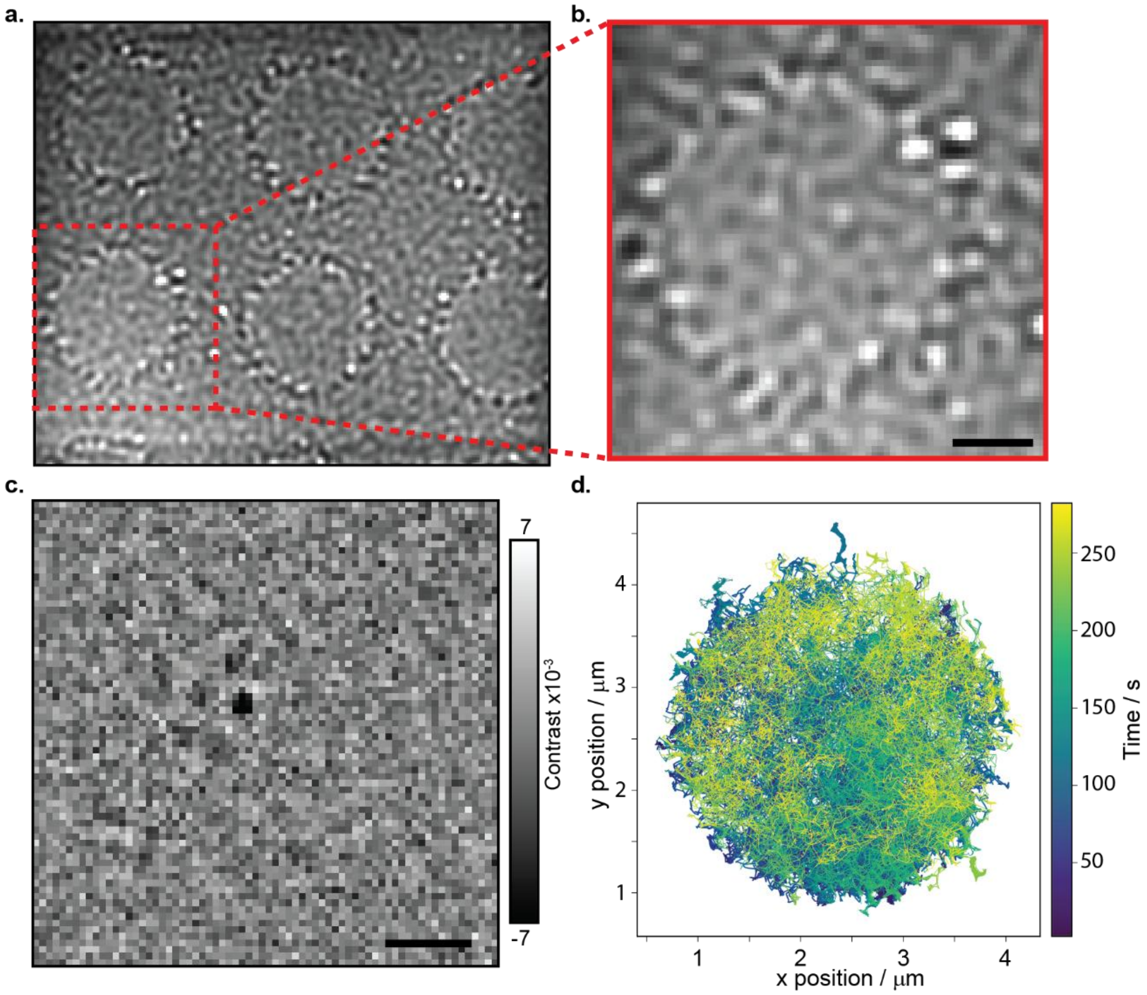

**Figure S20: Mass photometry  using confined SLBs** (a) MP image of the confined supported lipid bilayer traps measured with a FOV of 15.4 × 13.2 μm$^2$. (b) MP image of an individual circular trap. Scale bar is 1 μm. (c) Median processed MP image of the same region, showing a diffusing mi3 pentamer of trimers in its centre. (d) Trace of the *x,y* positions of a single diffusing assembling complex during 5 min. of measurement at 250 Hz.

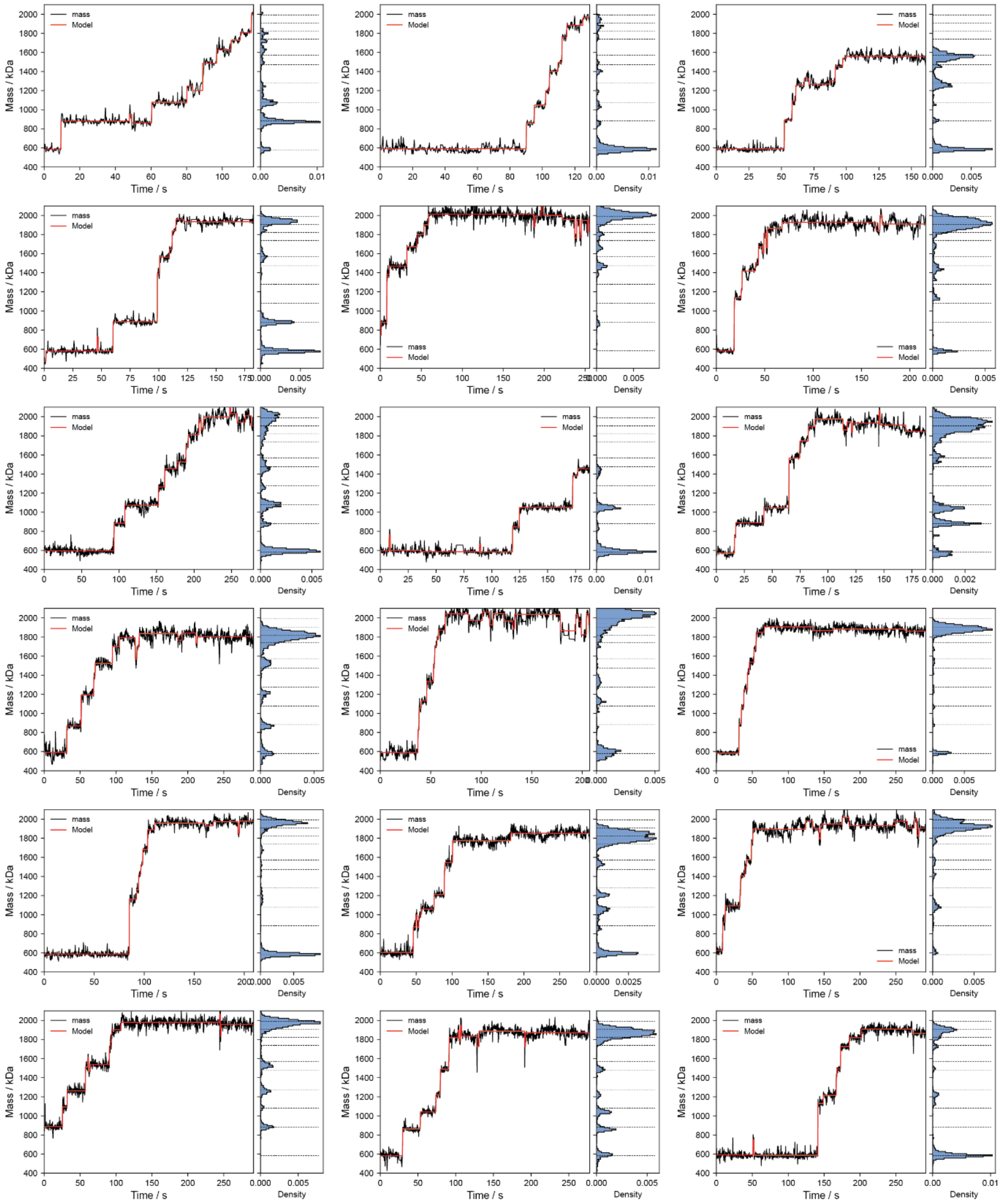

**Figure S21: Measured mass traces of the assembly process of mi3-VLPs in confined SLBs.** Each mass trace corresponds to a single VLP assembly process and contains: the measured mass trace as a function of time (black curve) following a moving median operation with a symmetric window size of 50 frames (200 ms) from each side, a step detection model (**Supplementary note 4.3**) that identifies transitions between oligomeric states (red curve), and a projected mass histogram. Dashed lines in the histogram, indicate the expected massed of the topologically closed states.

#### Supplementary movies captions

**Movie S1: Two-dimensional self-assembly dynamics of mi3-VLP subunits on a supported lipid bilayer.**

The left panel shows 5 s of a representative median-subtracted movie from a mass photometry measurement of mi3-subunits tethered to a supported lipid bilayer undergoing self-assembly. The movie was acquired at 270 Hz. The middle panel displays the raw mass-versus-time trace, with horizontal lines indicating the expected masses of the different oligomeric states. The right panel shows the oligomeric structures that correspond to the measured masses.

**Movie S2: Assembly of a pentameric ring into a complete mi3-VLP monitored by continuous mass photometry in molecular trap.**

The left panel shows 250 s of a representative median-subtracted movie from a mass photometry measurement of a pentameric ring tethered to a supported lipid bilayer. Upon addition of mi3-subunits to the solution, the pentamer assembles into a complete VLP. The middle panel displays the corresponding mass-versus-time trace, with grey dots indicating raw data at the native frame rate (250 Hz) and a black line representing a running median over 100 frames (400 ms). Horizontal lines mark the expected oligomeric masses of assembly intermediates, with red lines highlighting the masses of topologically-closed states. The right panel illustrates the structures of these topologically-closed intermediates along the assembly pathway.
